# Proto-sex locus in large yellow croaker provides insights into early evolution of the sex chromosome

**DOI:** 10.1101/2020.06.23.166249

**Authors:** Zhiyong Wang, Shijun Xiao, Mingyi Cai, Zhaofang Han, Wanbo Li, Yangjie Xie, Guicai Xu, Aiqiang Lin, Yan Zhang, Kun Ye, Huiyu Luo, Mengxiang Liao, Fang Han, Xiande Liu, Dongling Zhang, Qiurong Wang, Bi Wang, Weiming Li, Jiong-Tang Li

## Abstract

Autosomal origins of heterogametic sex chromosomes have been inferred frequently from suppressed recombination and gene degeneration manifested in incompletely differentiated sex chromosomes. However, the initial transition of an autosome region to a proto-sex locus has been not explored in depth. By assembling and analyzing a chromosome-level draft genome, we found a recent (evolved 0.26 million years ago), highly homologous, and *dmrt1* containing sex-determination locus with slightly reduced recombination in large yellow croaker (*Larimichthys crocea*), a teleost species with genetic sex determination (GSD) and with undifferentiated sex chromosomes. We observed genomic homology and polymorphic segregation of the proto-sex locus between sexes. Expression of *dmrt1* showed a stepwise increase in the development of testis, but not in the ovary. We infer that the inception of the proto-sex locus involves a few divergences in nucleotide sequences and slight suppression of recombination in an autosome region. In androgen-induced sex reversal of genetic females, in addition to *dmrt1*, genes in the conserved *dmrt1* cluster, and the rest of the sex determination network were activated. We provided evidence that broad functional links were shared by genetic sex determination and environmental sex reversal.

## Introduction

In animals, sex determination strategies are either genetic (GSD) or environmental (ESD). In GSD systems, sex specific alleles or genes drive sex determination. Heterogametic sex chromosomes, including the XY, ZW, XO, and ZO chromosome pairs (Vallender and Lahn 2006), epitomize the GSD system and have been intensively studied. These highly differentiated sex chromosomes are thought to have originated from autosomes (Muller 1914; Livernois et al. 2012), largely based on evidence from studies of species with incompletely differentiated sex chromosomes harboring gender-specific loci that determine sexes. Many examples of intermediate stages of sex chromosome evolution have been documented in fish species. In medaka (*Oryzias latipes), dmy*, a duplicate copy of *dmrt1*, is a Y-specific DM-domain gene required for testis development (Matsuda et al. 2002). In Nile tilapia (*Oreochromis niloticus*), *amhy*, a Y-specific duplicate of the anti-Müllerian hormone (*amh*) gene, induces testis development (Li et al. 2015). However, empirical evidence for initial transition from autosomes to incompletely differentiated sex chromosome is limited, because species in the early phase of sex chromosome differentiation have not been identified yet. This transition is posited to be initiated by a heterozygous state in one autosome, suppression of recombination between homologous chromosomes, and degeneration of a sex-linked locus or loci (Rice 1987; Charlesworth 1991; Peichel et al. 2004). If there exists a species that is in possession of proto-sex chromosomes and also retaining ancestral teleost karyotypes, it will help us to validate the hypothesized autosomal origin of the sex chromosome and study the initial transition process and evolution of the sex chromosome.

In some GSD species, including insects (Narita et al. 2007), fish (Piferrer 2001), and bird (Smith et al. 2003), environmental sex reversal (ESR) alters the primary sex into the opposite sex without modifying the genotype (Stelkens and Wedekind 2010). The ESR reversal is widely used in aquaculture by exposing fish to environmental factors to produce the preferred sex (Stelkens and Wedekind 2010). Several genes have been reported to participate in both ESR and GSD (Smith et al. 2003; Kohno et al. 2015), suggesting that ESR and GSD share similar functional components and possibly the same origin (Crews and Bull 2009). We further speculate that in early evolution of sex determination mechanisms, GSD and ESR may have sequestered a similar group of genes to drive the eventual differentiation of gonads. If there exists a species in an early phase of sex chromosome evolution and having both GSD and ESR pathways, it would be ideal to define the genetic and functional link between the two sex determination mechanisms.

In this study, large yellow croaker (*Larimichthys crocea*, designated as ‘croaker’ hereafter), a species employing the XX/XY determination system (Ning et al. 2007), was found to maintain ancestral karyotypes of teleosts and proto-sex chromosomes. The special genomic features reveal the initial transition process from autosomes to sex chromosomes. We also took advantage of the hormone-dependent ESR that reverses females to phenotypic males (called as ‘pseudo-males’) in this species, and examined the molecular interactions between the GSD and ESR in the proto-sex chromosome.

## Results

### Chromosome-level assembly of the croaker genome

A chromosome-level genome assembly is useful to identify the sex-linked DNA sequences in species without heteromorphic sex chromosomes. The current croaker draft genomes from wild croakers were assembled at scaffold level (Wu et al. 2014; Ao et al. 2015), in which a high level of heterozygosity was expected. With a 200 x sequencing depth including reads from Illumina and Pacbio platforms, we assembled a new reference genome at the chromosome level from an adult female gynogen croaker (supplementary table S1). The genome generated by a hybrid strategy of assembly and gap closure had a size of 689 Mb, with a scaffold N50 of 1.23 Mb and a contig N50 of 406.4 kb (supplementary table S2). Over 90% of the assembly were composed of 626 scaffolds longer than 271 kb. The assembly covered 95% of the estimated croaker genome size, which was 725 Mb as estimated by the distribution of K-mer frequency (supplementary fig. S1) and flow cytometry (Chen Z 2014). The assembly quality metrics were better than those of previous two assemblies (supplementary table S2 and S3). To assemble the genome at the chromosome-level, we anchored scaffolds to a croaker genetic map (Xiao, Wang, et al. 2015), capturing 74% of genomic sequences in 24 linkage groups (supplementary fig. S2).

We further assessed the quality of the genome assembly with five indicators. Almost 90% of genomic sequencing reads were aligned to the assembly (supplementary table S4). Likewise, genome re-sequencing reads from four individual croakers and the two populations mapped to the assembly at an average ratio of 92.4% (supplementary table S5). Moreover, RNA-seq reads from multiple tissues (Xiao, Han, et al. 2015) had an average alignment ratio of 92% to the assembly using TopHat (Trapnell et al. 2009) and almost all proteins in CEGMA (Parra et al. 2007) were covered by the assembly (supplementary table S6). The insert size distributions of the five paired-end/mate-pair libraries were consistent with the estimated insert sizes (supplementary fig. S3). These data indicated that our chromosome-level assembly was highly accurate.

We predicted a consensus gene set containing 23,272 protein-coding genes (supplementary table S7) and confirmed their accuracy. Firstly, the gene structure metrics (protein length, exon number and exon length) were comparable to those predicted from other model teleost genomes (supplementary fig. S4 and table S8). Secondly, we found many redundant pairs in the gene sets of the previous two assemblies but not in the new gene set (supplementary table S9), consistent with the expectation that high-heterozygosity assemblies might contain redundant genes. Thirdly, compared to those from the previous two croaker assemblies, the number of genes having functional annotations from the new assembly is more consistent with those annotated from other teleost genomes. Homologue searches against UniProt database (Consortium 2015) and NR database showed that most croaker genes from the new assembly had orthologues to the genes annotated in other species (supplementary table S10). Finally, twenty-four new genes not predicted in the previous assemblies (Wu et al. 2014; Ao et al. 2015) were all successfully amplified by PCR (supplementary fig. S5 and table S11). Among the identified protein-coding genes, 18,470 genes were annotated to have at least one Gene Ontology (GO) term and 12,010 genes were mapped to 331 Kyoto Encyclopedia of Genes and Genomes (KEGG) pathways. We considered the new reference gene set to be of high quality and useful for follow-up analyses.

### Maintenance of ancestral karyotypes in the croaker genome

After speciation from the teleost ancestor, many fishes underwent genome re-arrangement (Howe et al. 2013; Wang et al. 2015; Liu et al. 2016), suggesting that the modern sex chromosome might be homologous to multiple autosomes, thus hampering the study regarding origin of the initial autosomal sex chromosome. A fish species that has both GSD and conserved karyotypes of the ancestor species might be useful to infer autosomal origin of sex chromosomes. We determined the phylogenetic distance between the croaker and the other teleosts, especially medaka. Medaka is thought to have preserved the teleost ancestral karyotypes from more than 300 million years ago (mya) (Kasahara et al. 2007), possesses incompletely differentiated sex chromosomes (Kondo et al. 2006) and gender-specific loci (Matsuda et al. 2002). We assigned 18,979 croaker gene models to 6,960 TreeFam functional families. The genes from the other nine vertebrate genomes, including seven teleosts, chicken (*Gallus gallus*), and human (*Homo sapiens*), were also assigned to TreeFam families. The number of croaker genes assigned to functional families was comparable with the annotations of other teleost genomes (supplementary table S12 and S13). The orthologue profiles of the five related teleosts, including croaker, stickleback (*Gasterosteus aculeatus*), European sea bass (*Dicentrarchus labrax*), medaka, and tetraodon (*Tetraodon nigroviridiś*) (supplementary fig. S6) revealed that a high proportion of gene families are common across teleosts. The phylogenetic tree based on 877 strict 1:1 orthologues from the ten analyzed genomes grouped medaka, European sea bass, stickleback and croaker together (fig. 1a). The separation between croaker and medaka was estimated to occur 250 mya. The distribution of synonymous substitution rates (*Ks* values) among 877 orthologous pairs between medaka and croaker had a mean value of 1.32 (fig. 1b). Therefore, we estimated a mean rate of 0.0053 substitutions per synonymous site per million year.

**FIG. 1.**
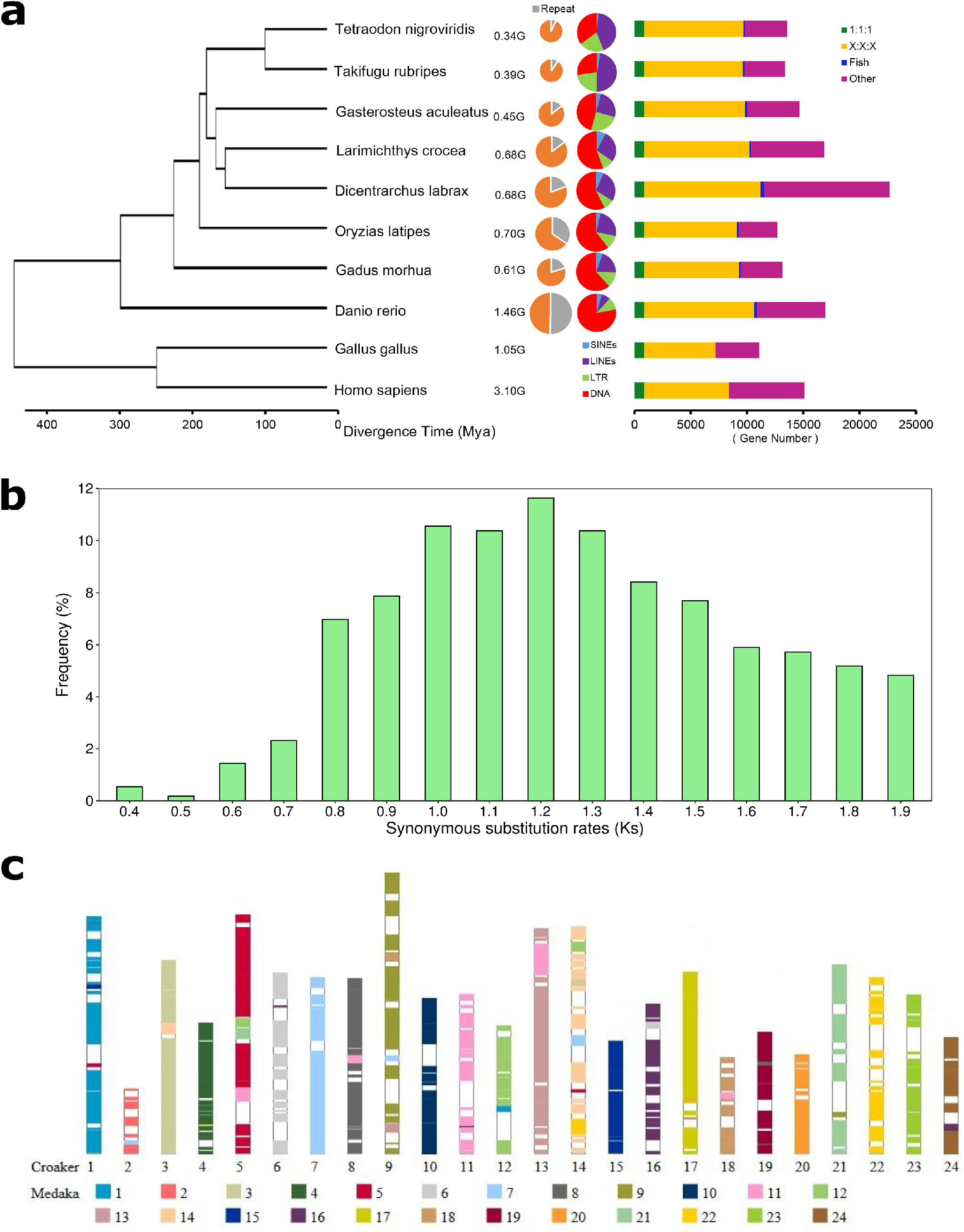
Comparative teleost genome analysis. (a) Phylogenetic tree, repeat components, and numbers of gene families among teleosts. Mya, million years ago. The grey part in the pie chart represented the proportion of repeat in each genome. Four types of color showed the proportions of four types of repeats out of all repeats. (b) The distribution of the synonymous substitution rates (*K*s) of strict 1:1 orthologue gene pair between medaka and croaker. (c) The distribution of the syntenic blocks in croaker genome. The syntenic blocks were represented by the colored bar with 24 codings based on their distributions on 24 medaka chromosomes.

Previously, a high-density genetic map has indicated that the croaker genome, with 24 chromosomes, has conserved synteny to the medaka genome (Xiao, Wang, et al. 2015). We speculated that croaker also maintains ancestral karyotypes as medaka. To test this hypothesis, we performed four-way chromosome-level synteny comparisons among the croaker, European sea bass, stickleback, and medaka genomes. Using the medaka genome as the reference ancestor chromosomes (Kasahara et al. 2007), we obtained 562 syntenic blocks with 6,597 orthologous gene pairs between the croaker genome and the medaka genome. The genome-wide comparison between medaka and croaker exhibited only minor lineage-specific chromosomal fusion events in croaker (fig. 1c). These results indicated that medaka and croaker had virtually identical distributions of syntenic blocks in their chromosomes, suggesting strongly that croaker, like medaka, also preserved the ancestral karyotypes in its genome.

We further speculated that croaker genome (725 Mb) and medaka genome (700 Mb) had similar proportions of gene components and transposable elements. Because the exonic regions of protein-coding genes across teleost genomes are virtually identical (supplementary fig. S4 and table S8), the variation in gene numbers and non-exonic size expansions might reflect the corresponding genome divergence. We found that the genome size was correlated with its gene number (*r* = 0.66, single-tail Student’s *t*-test *p* value = 0.039; fig. 2a), with its intronic regions (*r*= 0.985, single-tail Student’s *t*-test *p* value = 8×10^−6^; fig. 2b), and with its total intergenic regions (*r* = 0.998, single-tail Student’s *t*-test *p* value = 2×10^−8^; supplementary fig. S7a).

**FIG. 2.**
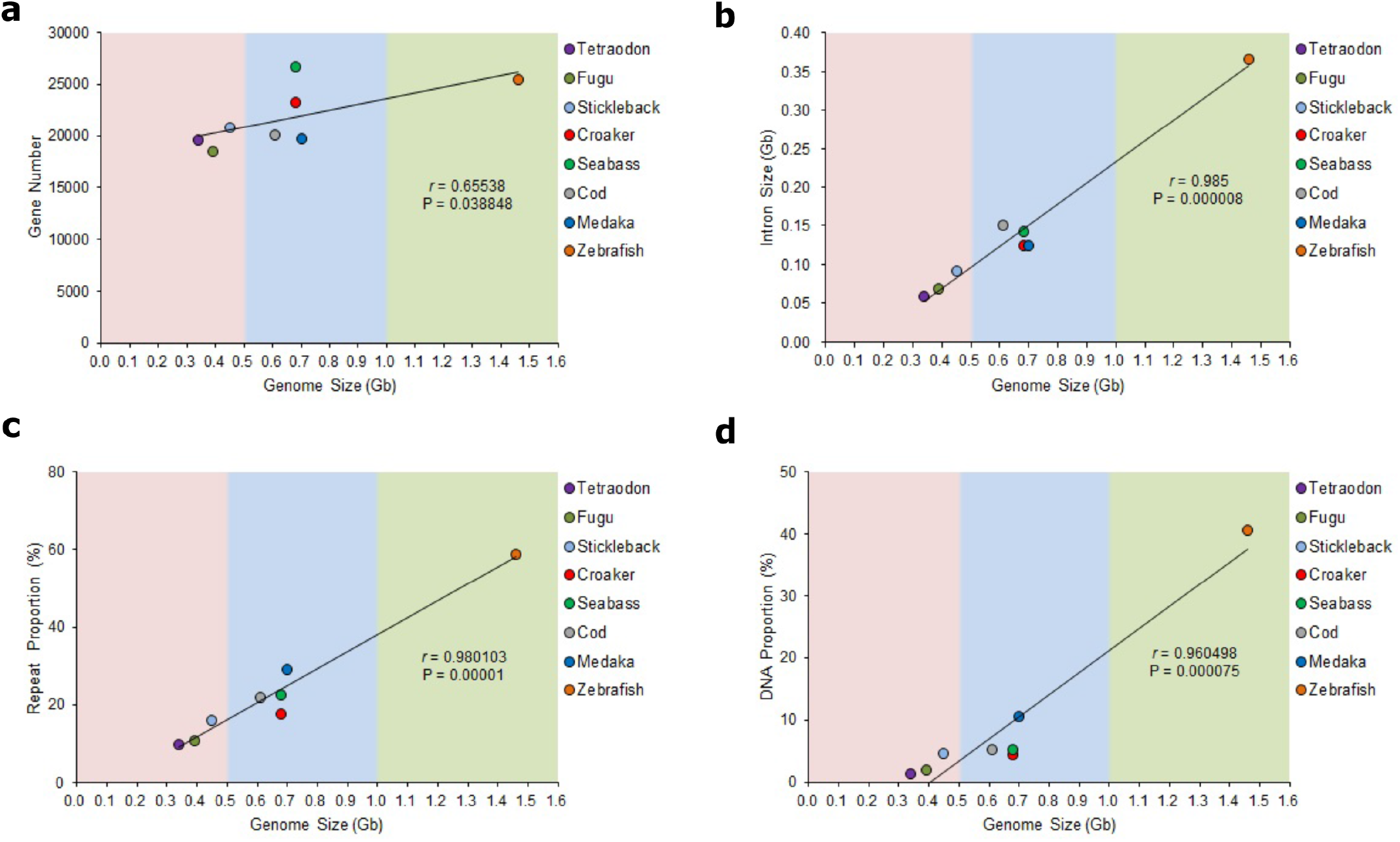
The expansion of teleost genome sizes. Teleost genomes were classified into three types: small-size genomes (pink), medium-size genomes (blue) and large-size genomes (green). (a) Significant correlation between gene number and genome size among eight teleosts. The dot with different color represented distinct species. (b) Intron size expansion in teleosts. (c) The correlation of repeat proportion and genome size. (d) The correlation of DNA transposon proportion and genome size.

We also suspected that croaker and medaka genomes contain a comparable proportion of repetitive elements. Repeat activity might drive genome size divergence. To study the proportions of repetitive elements among teleosts, we constructed *de novo* repeat libraries for croaker (supplementary table S14) and seven other teleosts (supplementary table S15). The croaker genome contained 518 consensus repetitive sequences, accounting for 14.6% of the assembly by *de novo* searches and homologue alignment. The classified repeat sequences accounted for 7.93% and the unclassified repeats comprised 6.67% of the croaker genome. Comparison among eight teleost genomes showed that the proportion of repeats was significantly correlated with genome sizes (*r* = 0.98, single-tail Student’s *t*-test *p* value = 1×10^−5^; fig. 2c), suggesting that the widespread repeat insertion may possibly have been a driving force for teleost genome expansion. We therefore examined the proportions of four types of repeat elements and their relationship to genome sizes in the teleosts studied. The proportions of DNA transposons was correlated with genome size (*r* = 0.96, single-tail Student’s *t*-test *p* value = 7.5×10^−5^, fig. 2d), so were those of short interspersed nuclear elements (SINEs) (*r* = 0.99, single-tail Student’s *t*-test *p* value = 1×10^−6^, supplementary fig. S7b) and long terminal repeats (LTRs) (*r* = 0.82, single-tail Student’s *t*-test *p* value = 6.65×10^−3^; supplementary fig. S7c). However, long interspersed nuclear element (LINE) proportions were not significantly correlated with genome size (*r* = 0.49, single-tail Student’s *t*-test *p* value = 0.11, supplementary fig. S7d). The activities of DNA transposons likely made more contributions to genome size expansions because these repeat elements accounted for higher proportions of genome content than other types of repeats. We clustered the teleost genomes into small, medium and large sized genomes based on their distributions of genes and transposable elements. The genomes in the same cluster showed similar gene numbers, gene sizes, and repeat proportions.

Taken together, comparisons between croaker and medaka genomes revealed phylogenetic closeness, virtually identical distributions of syntenic blocks, and proximate proportions of functional elements, providing strong evidence that croaker genome preserved the ancestor karyotypes.

### Male sex determination by a single proto-sex region

To identify the sex determination region in the croaker genome, we used a previously constructed and integrated genetic map (Xiao, Wang, et al. 2015) to perform QTL mapping and association study of sex in the same pedigreed family. The QTL mapping linked six markers in linkage group 9 (LG9) of the integrated map to the phenotypic sex under the LOD threshold of 30, ranging from 28 cM to 37.1 cM (fig. 3a and supplementary table S16). Subsequently, association studies confirmed that these six markers were associated with sex. Most of these markers exhibited heterozygosity in males and homozygosity in females (supplementary table S16). Based on the genotype difference in two sexes, we scanned genomic sequences of the six re-sequenced samples and identified 617 sites with polymorphism segregation around the GSD locus between the two sexes (supplementary table S17). These sites exhibited heterozygosity in males and homozygosity in females. The GSD locus was narrowed to a 4 Mb region on chromosome 9 (chr9), herein deemed the putative sex chromosome. This locus included 79 genes (supplementary table S18). Because croaker chr9 was homologous to medaka autosome 9 (fig. 1c), our results provide comparative genomics evidence that supports the hypothesis that sex chromosomes originated from autosomes.

**FIG. 3.**
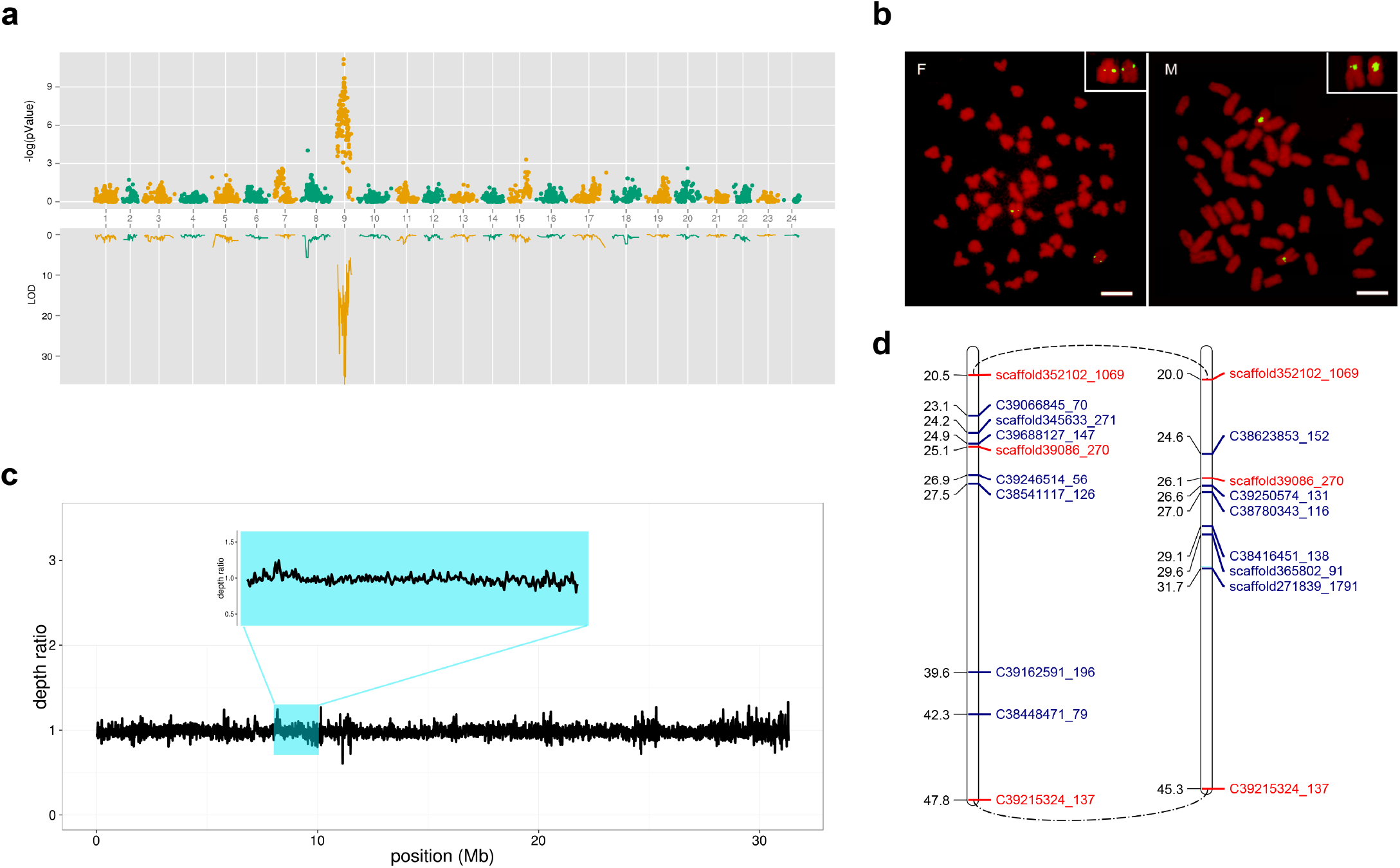
GSD locus in croaker. (a) GSD locus in chr9 (X axis) identified by the GWAS study (Y axis, top) and QTL analysis (Y axis, down). The significance of genetic markers (Y axis) was calculated in GWAS and LOD in QTL analysis. (d) BAC FISH analysis of croaker chromosomes showing a double signal in both males and females. A BAC clone containing the *dmrt1* gene, was labeled and used to probe chromosome spreads of male and female. Scale bars, 5 μm. (c) Relative sequencing depth (Y axis) between female population and male population. Relative sequencing depth ratio against the position along chr9 (X axis). The black lines represent the smooth ratio with a window of 1 kb. The small inserted figure illustrates the relative sequencing depth in the GSD region around *dmrt1*. (d) Slightly reduced recombination rate of the GSD locus in males. The blue markers are sex-linked in the female map (left) and male map (right). The consensus non-sex-linked markers in both maps are colored red. The arcs connect the markers around the GSD in the two maps.

The croaker GSD locus showed hallmark characteristics of sex determination regions of (incompletely) differential sex chromosomes. Firstly, this locus was significantly associated with the phenotypic sex. Secondly, some genes located in this region, or locus, were predicted to function in reproduction and male sex determination. Their orthologues, including *dmrt1* and *piwi*, have been shown to be associated with sex determination in other species (Megosh et al. 2006; Smith et al. 2009). Thirdly, this locus showed heterozygosity in males and homozygosity in females, as observed in both sex-linked markers and sex-biased variants. The significant segregation modes of the two variants in the two sexes were validated by genotyping another 15 male and 15 female croakers using Sanger sequencing (supplementary fig. S8 and table S19), demonstrating that these loci reliably identify gender in croaker. The status of allele segregation in the croaker GSD locus resembled those in (incompletely) differential sex chromosomes, where some loci were consistently heterozygous in one sex but homozygous in the other sex (Peichel et al. 2004). All three lines of evidence support the sex determination function of this locus in the croaker.

Notably, the putative sex chromosome (chr9) in croaker is homologous to medaka chromosome 9, the autosomal progenitor of the medaka sex chromosome (Kondo et al. 2006). Therefore, we posited that GSD locus in croaker chr9 may show characteristics of a proto-sex chromosome, which should be largely autosomal in nature (Spigler et al. 2010; Reisser et al. 2017). If this hypothesis is true, there should not be obviously differential sex chromosomes in croaker, as is expected in species with heteromorphic sex chromosomes (Chen et al. 2014). As expected, the estimated genome sizes of the two females and two males based on re-sequencing reads were almost equivalent, ranging from 726 to 730 Mb in both genders (supplementary fig. S9 and table S20). These re-sequencing reads from the males and females were aligned with the genome assembly at about the same ratios (Student’s *t*-test *p* value = 0.73, supplementary table S5). Moreover, FISH analysis with a BAC clone identified the putative sex chromosomes which were not visibly different cytogenetically in the males and female croakers (fig. 3b). These results were consistent with the prediction that croaker does not have obviously differential sex chromosomes.

If the GSD locus was under the early evolution of sex chromosome, the GSD divergence time between two genders should be very late, that is, the GSD was very young. We secondly estimate the GSD divergence time. Although 617 polymorphism sites within the GSD locus exhibited heterozygosity in males and homozygosity in females, only 13 sites were distributed in the coding regions of ten genes (supplementary table S21). We reconstructed protein sequences of male counterparts of these ten genes based on the polymorphism sites and revealed that the mean synonymous substitution rate (*Ks* value) of these genes between male genome and female genome was 0.0014. Assuming that the divergence rate between male and female GSD was equal to the rate of 0.0053 substitutions per synonymous site per million year, we estimated a mean GSD divergence time of ~0.26 mya between male and female.

For species with incompletely differentiated sex chromosomes, including medaka (Nanda et al. 2002), tilapia (Li et al. 2015) and stickleback (Peichel et al. 2004), the Y chromosomes have accumulated duplicates of sex-determination genes. Assuming that the croaker chr9 is a proto-sex chromosome, we thirdly predicted that there should not be duplicated sex-determination genes in the GSD locus in female or male genomes. We compared the relative sequencing depths of the GSD region in females and males, and found their ratio to be approximately 1 (fig. 3c). Moreover, FISH analyses showed two clear signals in both females and males (fig. 3b). This was different from the distinct signals in species with gene duplication or deletion in sex chromosome (Chen et al. 2014), and indicated a lack of gene duplication or loss in the GSD locus in both female and male croaker.

Our fourth prediction was that recombination is slightly reduced around the GSD locus in male croaker, but not suppressed or non-recombinant as shown in incompletely differential sex chromosomes or in intact sex chromosomes, respectively (Tennessen et al. 2016). The region around the GSD locus determined by two consensus non-sex-linked markers (*scaffold352_102 1069* and *C39215324_137*) was 27.3 cM in length in the female map and 25.3 cM in length in the male map, with a ratio of 1.08:1 (fig. 3d). This ratio was much lower than the sex specific recombination rate differences in nascent sex chromosomes (Peichel et al. 2004) or incompletely differential sex chromosomes (Singer et al. 2002; Phillips et al. 2009). These results suggested that recombination around the GSD locus was only slightly suppressed in male croakers. The results were also consistent with our re-sequencing results that croaker GSD were highly homologous between males and females (supplementary table S5), in contrast to the much lower level of homology in the incompletely differential sex chromosomes (Peichel et al. 2004; Takehana et al. 2014).

Our search for the sex chromosome yielded a single and young proto-sex region in chr9 that was clearly linked to male sex determination. The incompletely differential sex chromosomes lost the autosomal nature, having either duplication (loss), much reduced recombination rate or poor sequence homology. In contrast, the young croaker GSD region displayed largely autosomal features, including a lack of obvious differentiation, no gene duplication or loss, a slightly reduced recombination rate, and a high level of sequence homology. We therefore deemed that croaker chr9 is a proto-sex chromosome, which differs from the incompletely differential sex chromosomes.

### Differential expression and polymorphic segregation of *dmrt1* in male and female croaker

To infer how chr9, which is virtually an autosome, could determine sex, we examined whether candidate sex-determination genes in the GSD locus have polymorphisms that are perfectly segregated between sexes, in accordance to the zygosity difference of sex chromosomes. Among the 79 genes encoded in the GSD region, *dmrt1* displayed many features characteristic of a master sex-determining gene.

Firstly, *dmrt1* is functionally well-conserved. Its orthologues in chicken and many fishes participate in male sex determination (Smith et al. 2009; Chen et al. 2014). We identified six *dmrt* members in croaker, consistent with the six orthologues in other teleosts, while human has nine and chicken has four. Phylogenetic analysis revealed that seven sub-families (*dmrt1, dmrt2, dmrt3, dmrtA1, dmrtA2, dmrtB1* and *dmrtC*) of *dmrt* were present prior to the teleost/tetrapod divergence (supplementary fig. S10).

Secondly, the *dmrt1* cluster is well-preserved across teleosts and remains intact in croaker genome. A micro-syntenic cluster consisting of *dmrt1, dmrt3, dmrt2*, and nine neighboring genes, was highly conserved across teleosts. This cluster exhibits the same gene order across teleosts (fig. 4). The croaker *dmrt1* cluster resembles the intact genes of this conserved region, except a croaker-specific gene (*rnf*). These results suggested that the teleost ancestor might already have a 12-gene *dmrt1* cluster.

**FIG. 4.**
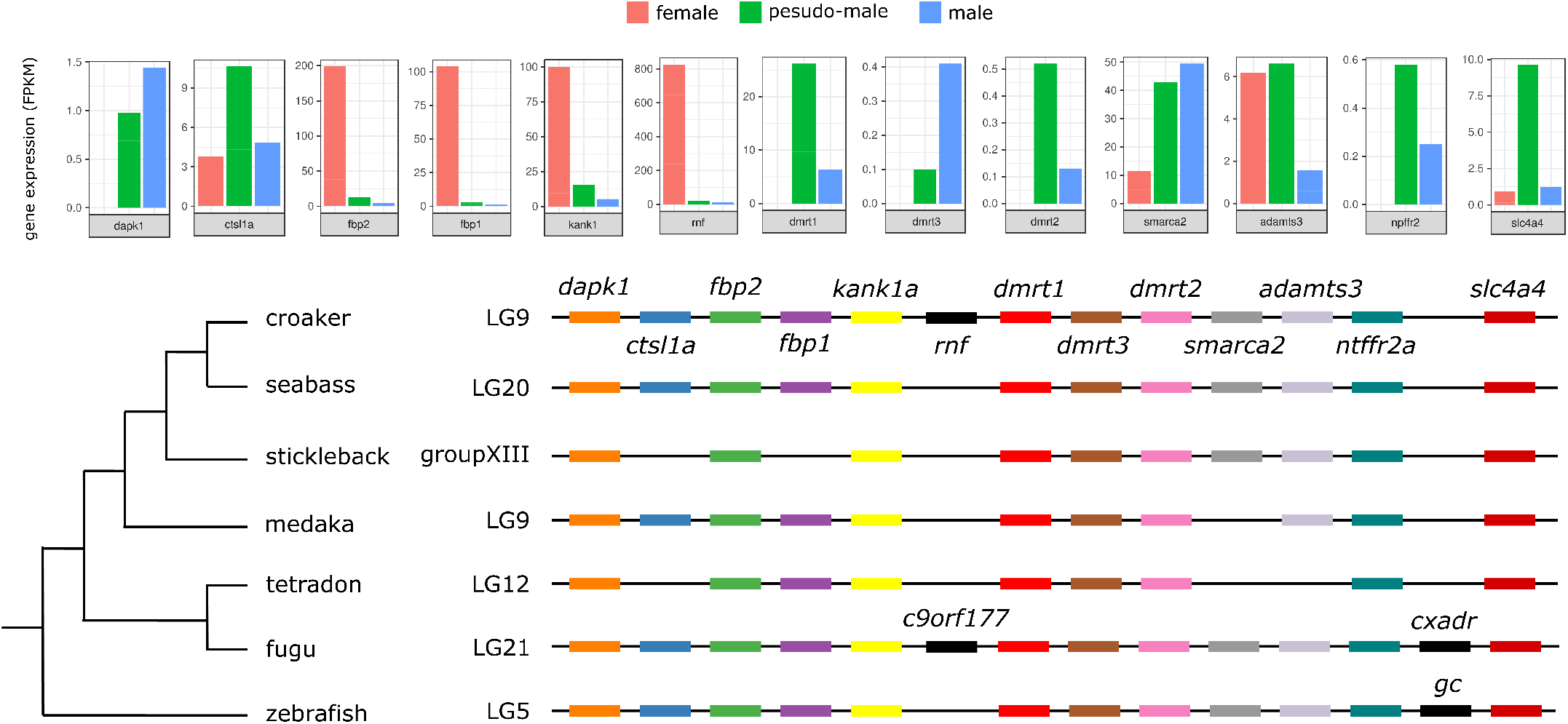
The conserved *dmrt1* gene cluster and the expression profiles in gonads of females, males, and pseudo-males. Chromosomal organization of *dmrt1* and neighboring genes of teleosts is displayed at the bottom. This cluster exhibits the same gene order and is present as intact across teleosts, possibly representing the ancestral state of these clusters in the teleost ancestor. The species-specific genes in this cluster are colored black. The FPKM values for gene expressions of the 13 genes in the *dmrt1* cluster in gonads of female (red), pseudo-male (green) and male (blue) are shown at the top. Four genes were up-regulated in female gonads, whereas expression of five genes had higher expression levels in pseudo-males and males.

Thirdly, croaker *dmrt1* exhibited male-specific expression in testis, as shown in RNA-seq and qRT-PCR analyses (supplementary fig. S11 and fig. S12). The testis expression of *dmrt1* increased stepwise at 6, 13, 20, 27, 34, 41, 49, 55, 69, 84, 96, 112, 123 days after hatching (dah), and in 8 months, 16 months, and 2 years old males, along with the maturation of testis (supplementary fig. S13). In particular, the transcriptional level of *dmrt1* increased progressively during the larval development in males, peaking at 112 dah, when distinct morphology between female and male gonads becomes apparent. Taken together, the sex-linkage, phylogenetic tree, male-specific expression of *dmrt1* that coincides with testis development implicated this gene as a critical regulator of male development in croaker.

Finally, we examined the segregation mode of polymorphism variants around *dmrt1*. Among the 483 SNPs in the region, starting from 3 kb upstream and ending at 3 kb downstream of *dmrt1*, 34 had significant segregation mode between males and females. These 34 SNPs were distributed in UTR regions, introns, and transcription binding sites (supplementary table S22). All exonic SNPs resided in the UTR regions but not in any of the miRNA target sites. Three SNPs were located at the binding sites of transcriptional factors including TATA-binding protein (TBP), Snail, Broad-complex_1, c-FOS, FREAC-2, and SQUA (supplementary fig. S14). Notably, TBP is known to regulate mRNA levels during testis development (Schmidt and Schibler 1997) and *c-FOS* expression is male-biased in zebrafish (*Danio rerio*), which is linked to sexual behavior (Pradhan and Olsson 2015). Since the only genetic difference in the *dmrt1* regions between the two genders was SNPs, these polymorphisms may be the causes for the segregation of *dmrt1*, but how such polymorphisms at the single nucleotide level lead to differential expression of *dmrt1* remains an open question.

### Extensive function link between GSD and hormone-induced ESR in croaker

We speculated that a direct comparison of gene networks induced by GSD and hormone-induced ESR would provide further information regarding GSD in croaker. Previous studies have shown that hormone-induced ESR and GSD may share similar functional components (Smith et al. 2003; Kohno et al. 2015) and possibly the same origin (Crews and Bull 2009). We therefore created pseudo-male croakers by treating the females with 17α-methyltestosterone. The ovaries of treated individuals were pervaded with spermatogonia, spermatocyte, sertoli cells and spermatozoon (fig. 5a), comparably as in testes, and in direct contrast to ovaries that mainly consisted of follicular epithelia, oogonia and oocytes. Through RNA-seq analysis of these three types of gonads (supplementary table S23), we identified 8,349; 7,664 and 2,929 differentially expressed genes (DEGs) in pairs between females vs males, females vs pseudo-males, and males vs pseudo-males, respectively (supplementary fig. S15). Sample clustering using the expression levels of all DEGs revealed that pseudo-males were closely to males, which was consistent with gonad histological analysis (fig. 5b).

**FIG. 5.**
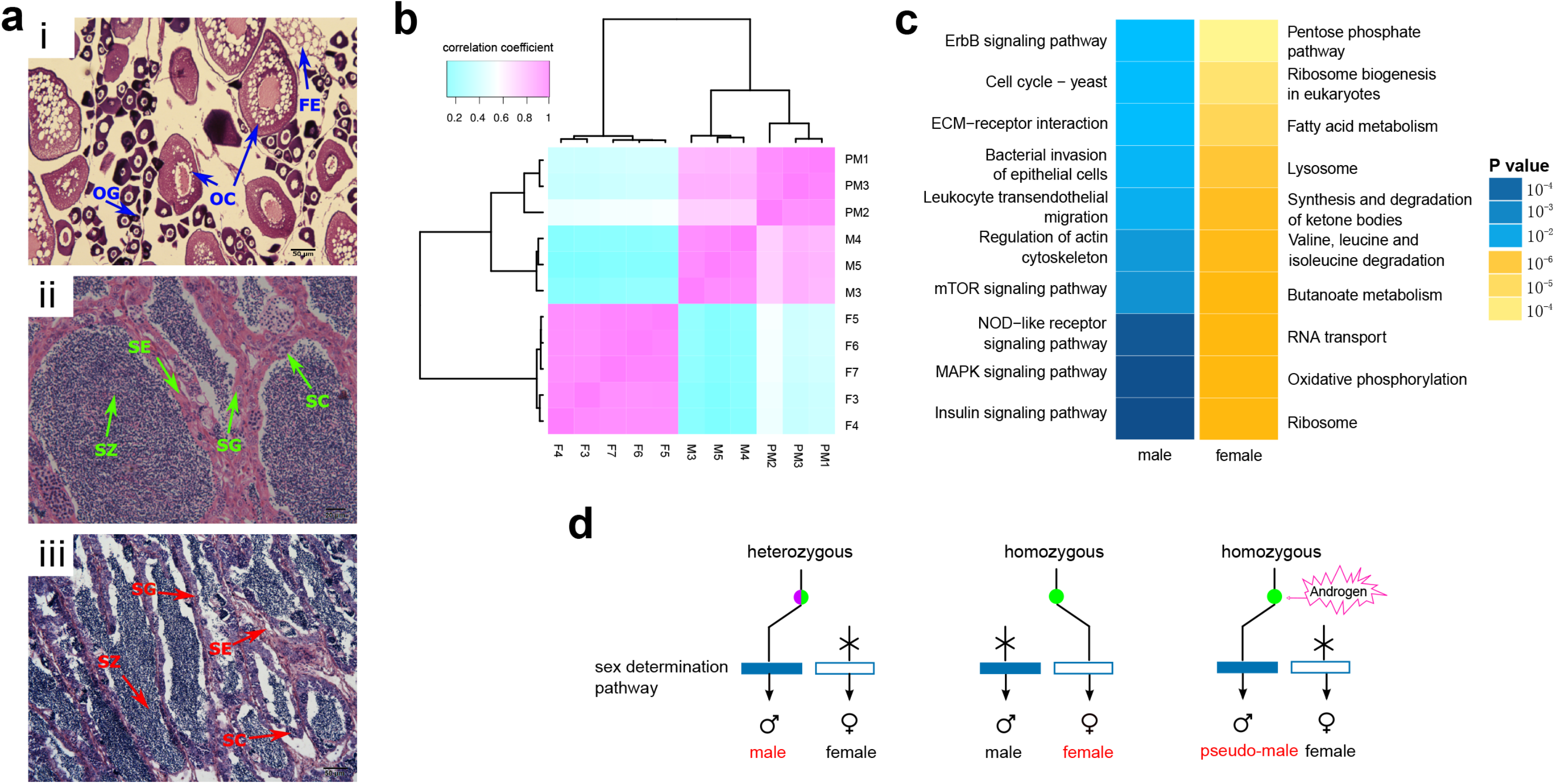
Characterization of hormone-induced ESR in croaker. (a) Morphological analysis of gonads in females, males and pseudo-males. Gonad histological observed for (i) females, (ii) males, and (iii) pseudo-males. FE, follicular epithelia; OG, oogonia; OC, oocyte; SG, spermatogonia; SC, spermacyte; SE, sertoli cell; SZ, spermatozoon. (b) Sample clustering of females, males and pseudo-males using the expression values of DEGs. The sample IDs were listed in supplementary table S23. (c) The enriched pathways by common DEGs shared in GSD and ESR. Blue represents pathways enriched in both males and pseudo-males and red represents pathways enriched in females. (d) A proposed mechanism of male determination co-operated by both GSD and ESR. The node (colored purple and green) represents heterozygous genotype and the green node indicates homozygous genotype in the GSD locus. The blue and white bars are the male and female sex determination pathways, respectively. Without the treatment of androgen, the male sex determination pathways were activated if the GSD region were heterozygous. The pathways were switched off when the homozygosity of GSD locus. If treated with androgen, the male sex determination pathways were activated even when the homozygosity of GSD region, generating pseudo-males.

Firstly, we examined shared gene expression patterns in the three gonad types to study gene components between sex determination and reversal. Using the DEG set between male and pseudo-male gonads as a negative control, we identified 6,316 hormone-induced DEGs (females vs pseudo-males), including 3,614 up-regulated genes (UEGs) in pseudo-males and 2,702 UEGs in females. Through comparison of females and males, we identified 6,344 genetic-determining DEGs, which consisted of 3,241 UEGs in males and 3,103 UEGs in females. Comparison of hormone-induced DEGs and genetic-determining DEGs revealed 4,674 common DEGs in GSD and in hormone-induced ESR, including 2,289 UEGs in females and 2,385 UEGs in both males and pseudo-males (supplementary fig. S15). These UEGs might be core gene components shared by GSD and ESR. They accounted for 73.7% of GSD DEGs and 74% of ESR DEGs. These results revealed that a substantive number of genes are shared by ESR and GSD.

Secondly, we examined whether GSD genes had differentially expression profiles among the three types of gonads. A total of 18 GSD genes were consistently up-regulated in both pseudo-males and males, and 13 GSD genes had higher expression levels in females than in pseudo-males and males (supplementary table S24). Interestingly, among the genes in the *dmrt1* cluster, the expressions of *fbp2, fbp1, kank1*, and *rnf*, were up-regulated in females. In contrast, the genes of *dmrt1, dmrt3, dmrt2, smarca2*, and *npffr2*, had higher expression levels in pseudo-males and males (fig. 4). The qRT-PCR analysis confirmed the differential expressions of two GSD genes (*dmrt1* and *rnf*) among females, males and pseudo-males (supplementary fig. S12 and table S25). The result suggested that some of GSD genes, including genes in the *dmrt1* cluster, might participate in the regulation of sex-reversal triggered by hormones.

*Dmrt1* is a master sex-determining gene and exhibits consistent expression patterns in both GSD and hormone-induced ESR. Thus, to understand the gene regulatory network including *dmrt1*, thirdly we examined genes having highly similar expression profiles with *dmrt1*. Totally 66 genes were identified to have a close expression relationship with *dmrt1* among three types of gonads using weighted gene correlation network analysis (WGCNA) (Langfelder and Horvath 2008) with a weight threshold of 0.7 (supplementary fig. S16, and table S26). Functionally, the involvement of *tesk1* (Toshima et al. 1995; Otani et al. 2017) and *zp3* (Cui et al. 1994; Hirakawa et al. 2012) has been previously reported to participate in both sex determination and sex reversal. This result further revealed that besides GSD genes, significantly co-expressed genes with *dmrt1*, might participate in the regulation of hormone-induced sex-reversal.

Finally, we analyzed the global functional links between GSD and ESR. In the comparative analysis of females and pseudo-males, the pseudo-male UEGs were enriched in ECM-receptor interaction, MAPK signaling pathway, insulin signaling pathway and other signaling pathways (50 pathways), some of which had been demonstrated to be crucial to male sex determination (Nef et al. 2003; Bogani et al. 2009) (supplementary table S27). The female UEGs were enriched in oxidative phosphorylation, steroid hormone biosynthesis, and other processes (63 pathways). In the comparative analysis of females and males, the male UEGs were enriched in male-determination related pathways, including insulin signaling pathway, NOD-like receptor signaling pathway, and MAPK signaling pathway (42 pathways, supplementary table S28). The pathways enriched by female UEGs in this comparison (69 pathways) were similar to those pathways enriched by female UEGs as demonstrated in the previous comparative study between pseudo-males and females. The shared 2,289 female UEGs by ESR and GSD were involved in 50 pathways, including oxidative phosphorylation and steroid hormone biosynthesis (fig. 5c and supplementary table S29). The shared 2,385 UEGs in both males and pseudo-males were enriched in male-determination related pathways (31 pathways, fig. 5c and supplementary table S29). The shared pathways (81) accounted for 71.7% of ESR pathways (113) and 73.0% of GSD pathways (111). The results indicated that GSD and hormone-induced ESR likely share similar biological pathways.

The comparisons revealed that ESR and GSD had extensive functional links that shared similar gene components and biological pathways, rendering further support to the hypothesis that they might be of the same origin. Since that croaker GSD resides in a pair of autosomes that is at a much early stage of differentiation to sex chromosomes, we hypothesized that ESR mechanism might have evolved since the phase of autosome differentiation.

## Discussion

We have found a *dmrt1* based GSD locus in an autosome that appears to manifest the initial transition from the autosome to sex chromosome. Our genomic and genetic data revealed that this locus exhibited features of genetic sex determination: (1) significant association with phenotypic sex; (2) orthologues of some genes functioning in reproduction and male sex determination; and (3) sequence heterozygosity in male and homozygosity in female. Our results further revealed autosomal characteristics of the GSD locus: (1) no apparent difference between sexes; (2) no duplication or loss of genes; (3) slightly reduced recombination rate in the male sex; and (4) high sequence homology between the male and female genome. We also found that the GSD locus is very young. We conclude that croaker chr9 is reminiscent of a much earlier phase of sex chromosome evolution than that of incompletely differential sex chromosomes with gender-specific loci.

The maintenance of the teleost ancestral karyotype helps us to trace the evolution of the GSD region from teleost ancestor to modern fish. Brunner *et al*. identified a three-gene cluster consisting of *dmrt1, dmrt2*, and *dmrt3*, highly conserved from fish to mammals (Brunner et al. 2001). We further found that this highly conserved cluster was extended to a 12-gene region in teleost. A *dmrt1* enabled SD mechanism has been confirmed in many animal taxa (Ferguson-Smith 2007; Hong et al. 2007; Smith et al. 2009; Chen et al. 2014). We also demonstrated that this highly conserved region is potentially a GSD locus in croaker. Taken together, this micro-syntenic cluster might already exist as a GSD locus in teleost ancestor. In modern fish, this ancestral GSD locus has been transferred to different chromosomes. In croaker, this locus still locates on the autosome 9; while in medaka, it underwent a duplication event and the duplicated locus finally harboured on the nascent Y chromosome (Kondo et al. 2006).

This nascent GSD locus supports the hypothesis that heterogametic sex chromosomes originated from autosomes. It also provides us the opportunity to study the contributions of different factors to suppress the recombination. Firstly, sequence expansion by repeat accumulation, gene duplication or acquisition of additional genes might result in suppression of recombination in the sex determination region (Bergero et al. 2013; Murata et al. 2015). However, our results show slightly reduced recombination suppression without any expansion, acquisition, or loss in the proto-sex locus between two genders (fig. 3c). Secondly, another explanation for recombination suppression is sexual antagonism (Jordan and Charlesworth 2012). Sexual polymorphism selects for reduced recombination with the GSD region, increasing transmission of the benefit alleles to one gender and inhibiting transmission of these alleles to the other gender (Bergero et al. 2013). Indeed, we observed that mild recombination suppression was coupled with sex-segregated SNPs. Thirdly, chromosome rearrangement, including inversion and conversion, is supposed to mediate suppression of recombination (Fraser and Heitman 2005). We aligned the previously assembled draft male genome sequences (Xiao, Wang, et al. 2015) to the reference assembly with BLAT. If one sequence from male assembly was broken into two discontinuous and intra-chromosomal regions in the reference assembly, we considered that there existed a rearrangement event. No rearrangement in the male assembly was observed compared to the reference assembly. One possible explanation is that chromosome rearrangement has not occurred in the nascent GSD locus yet (Murata et al. 2015). The other explanation is that it may be due to the low-quality male genome assembly. A chromosome-level male genome assembly is required to examine whether rearrangement exist in the GSD locus of one gender. Based on this nascent GSD model, we speculate that the initial recombination suppression following sex chromosome differentiation should be initially mediated by sexual antagonism, which was at least prior to sequence expansion.

Our comprehensive investigation of gene expression in male, female and pseudo-male croakers revealed extensive functional links of regulated genes and biological processes between GSD and hormone-induced ESR, suggesting that these two mechanisms might have a same origin. Because the proto-sex chromosome was preserved in croaker, ESR mechanism might have evolved since autosome differentiation and the link between GSD and ESR was ancient. On the basis of the phenotype consistency and high overlapping of DEGs, we propose a switch-like sex determination model in croaker (fig. 5d). Genetic zygosity functions as the primary mechanism that controls sex determination where heterozygous genotype determines male phenotype. In contrast, the homozygous genotype results in female phenotype, which can be switched to the male phenotype by the androgen treatment. The genetic-driven and hormone-induced male phenotypes appear to recruit a similar set of genes to determine sex. This proposed mechanism might improve our knowledge related to the interaction between two different pathways.

## Materials and Methods

### Sequencing on Illumina and Pacbio platforms

A gynogenetic croaker was generated (Komen and Thorgaard 2007) and selected as the genomic DNA source for whole-genome sequencing (supplementary note 1) to decrease interference of genomic heterozygosity to assembling. Two pair-end libraries and three mate-pair libraries (supplementary table S1) were sequenced on the Illumina platform. The genomic DNA molecules were also sequenced using the PacBio Sequel platform. Two 20 kb libraries were constructed for the Sequel platform, and about 11.2 Gb long reads with N50 length of 13.7 kb were generated (supplementary note 2).

Secondly, we randomly selected four individuals (two females and two males) for whole genome re-sequencing and constructed one paired-end Illumina library for each individual. Thirdly, we extracted the genomic DNA of another fifty females and fifty males and pooled the DNA of samples of the same gender together (supplementary note 1). For each population, we constructed one pair-end Illumina library and sequenced it with 2 × 100 bp using Illumina platform. The re-sequencing reads were cleaned using SolexaQA (Cox et al. 2010). The re-sequencing data were listed in supplementary table S5.

Fourthly, we prepared pseudo-males by treating females with 17α-methyltestosterone (supplementary note 3). We performed RNA-seq sequencing on the gonads of two females, three males and three pseudo-males (supplementary note 4). The sequencing for samples of each gender was run in triplicate with 100 PE mode (supplementary table S23) on the Illumina platform. The RNA-seq reads were trimmed to filter out low-quality bases.

### Construction of chromosome-level genome assembly

We filtered out low-quality and short reads to obtain a set of usable reads using SolexaQA (Cox et al. 2010). The raw Pacbio reads were corrected using MECAT (Xiao et al. 2017). Before assembly, we estimated the genome size of female croaker using Jellyfish (Marçais and Kingsford 2011) (supplementary note 5). The cleaned Illumina reads were assembled and further scaffolded. The gaps in these scaffolds were closed using the pair-end reads and error-corrected long reads, respectively (supplementary note 6). Finally, the closed assembly was anchored to a high-dense genetic map of croaker consisting of 3,448 markers in 24 linkage groups (Xiao, Wang, et al. 2015).

### Gene prediction and function annotation

For the *de novo* gene prediction, RepeatModeler (http://www.repeatmasker.org/RepeatModeler/) was used to construct a croaker specific repeat library. Using RepeatMasker (http://repeatmasker.org/), the genome assembly was masked firstly with this specific library and secondly with teleost repeat library in RepeatMasker package.

We predicted protein-coding genes in the repeat-masked assembly using a combination of *de novo* predicted genes, homology-based predicted genes, and RNA-seq based models (supplementary note 7). The cuffmerge package in Cufflinks (Ghosh and Chan 2016) was used to combine the three sets of predicted gene models and generate a consensus gene set. In each consensus gene, the transcript model having the longest CDS was selected to represent the gene model.

We then annotated the functions for croaker protein-coding genes. Homologues were identified using Blastp (e value of 10^−5^) against Swiss-Prot, TrEMBL (Consortium 2015) and NCBI non-redundant (NR) protein database. The KEGG biological pathways to croaker genes were annotated using KEGG Automatic Annotation Server (KAAS) (Moriya et al. 2007). For Gene Ontology (GO) analysis, we used Blast2GO (Conesa et al. 2005) to annotate the GO information of croaker genes.

### Assessment of the qualities of genome assembly and gene prediction

The quality of the assembly was evaluated using the metrics described below: (1) the proportion of *de novo* genome sequencing reads aligned to the assembly by BWA (Li and Durbin 2010); (2) the insert size distributions of paired-end/mate-pair libraries; (3) the mapping ratio of RNA-sequencing reads to genome using bowtie2 (Langmead and Salzberg 2012); (4) CEGMA-based evaluation by aligning 248 core eukaryotic genes dataset (CEGs) to the assembly with BLAT (Kent 2002); and (5) the mapping ratio of genome re-sequencing reads by BWA (Li and Durbin 2010).

We further assessed the quality of the predicted genes using the metrics described below: (1) comparing gene structures of croaker against those of other model teleost; (2) examining whether there exist identical predicted genes; and (3) validation of the predicted genes by amplifying 24 selected genes (supplementary note 8).

### Estimation of repeat components among teleosts

To study the repeat components across teleosts, the assembled croaker genome and other teleost genomes were masked as follows: (1) Using RepeatScout (Price et al. 2005), the consensus sequences of repetitive families in each species were generated and classified into four categories according to their mechanisms of transposition (including DNA transposons, LTRs, LINEs and SINEs). To search the *de novo* repeats in the genome, the genome was searched against the *de novo* repeat libraries using RepeatMasker with the default parameters; (2) After masking the genome using the *de novo* repeat library, each genome was further searched for homologous repeats using RepeatMasker against RepBase (Bao et al. 2015) fish library.

We calculated the total repeat proportion and the proportions of four categories described above in each genome by combining the *de novo* predicted repeats and homologous repeats. The correlation between repeat proportion and genome size was measured with Pearson’s correlation.

### Phylogenetic analysis and synteny comparison

Phylogenetic analysis was performed among ten species, including stickleback, medaka, takifugu (*Takifugu rubripeś*), tetraodon, croaker, European sea bass, Atlantic cod (*Gadus morhua*), zebrafish, chicken, and human. With the exception of European sea bass (downloaded from sea bass genome database: http://seabass.mpipz.mpg.de), proteins were obtained from Ensembl (Aken et al. 2017). We followed the strategy of Wang *et al*. (Wang et al. 2015) to assign genes of the ten species to TreeFam (Schreiber et al. 2014) gene families by the Treefam_scan with the cut-off of 1×e^−10^ and score over 100. Those families with one gene per species were used to construct a phylogenetic tree for the ten genomes. The proteins in each family were aligned with MUSCLE (Edgar 2004) and then the protein alignments were converted to codon alignments with PAL2NAL (Suyama et al. 2006). Gblocks (Talavera and Castresana 2007) was used to refine each codon alignment. All refined codon alignments were concatenated to form one super-alignment. PhyML (Guindon et al. 2005) was used to reconstruct the phylogenetic tree using the best nucleotide substitution model (GTR+gamma+I) of the super-alignment. Human was assigned as the out-group of the phylogenetic tree. The divergence time for croaker and other teleosts was estimated using the calibration time of human and zebrafish (429 mya), obtained from the TimeTree database (Hedges et al. 2006).

To estimate the rate of substitutions per synonymous site per year, two sequences in each 1:1 orthologue pair between medaka and croaker were aligned with ClustalW (Larkin et al. 2007) and the synonymous substitution rates of the corresponding coding sequences were calculated using the CodeML program from the PAML package (Yang 2007). We calculated a mean rate of silent substitution per year based on the divergence time between medaka and croaker.

We selected croaker and other closely related fish (medaka, European sea bass and stickleback) to analyse synteny of ancestral teleost karotypes. All-against-all alignments among proteins from four species were performed using Blastp with an e-value cutoff of 1×e^−5^. Medaka genome, which maintains the teleost karyotypes, was selected as a reference. The other three species were compared to the medaka genome to identify syntenic blocks between each of two species using MCScanX (Wang et al. 2012). Two compared chromosome regions with the gap size set to 15 genes and at least five genes were considered to be syntenic. Each fish had species-specific gene duplications, which could result in one-to-many syntenies. Therefore, we retained the one-to-one syntenies to trace teleost karyotypes in each species

### Identification of GSD locus

To compare the genome sizes of females and males, we estimated the genome sizes of two females and two males with whole genome re-sequencing data using Jellyfish (Marçais and Kingsford 2011). To identify croaker GSD loci, we performed QTL mapping and association analysis for the sex on one croaker family based on the integrated high-density genetic map consisting of 3,448 markers (Xiao, Wang, et al. 2015). The sex of the offspring was checked by gonad examination. Based on this published high-density genetic map of croaker, MapQTL (Van Ooijen and Kyazma 2009) was used to identify sex linkage regions. Composite interval mapping and multiple QTL model (MQM) mapping with a LOD threshold of 30 were utilized to detect significant markers associated with the sex. An association study with the simple linear regression model was applied to identify loci associated with sex using PLINK package (Purcell et al. 2007). Markers with *p* value < 1 ×e^−5^ were considered to be significantly associated.

To examine whether there exist duplicated genomic regions in females or males, we compared the relative sequencing depths of the GSD region between females and males. The cleaned genomic sequencing reads of female individuals and pooled male individuals were mapped to croaker genome by BWA (Li and Durbin 2010), respectively. Then we calculate the sequencing depth of the GSD regions with a 1,000 bp sliding window. Read depths were normalized by the total sequencing bases of the male population and the female population, respectively. The relative sequencing depth of each sliding window was calculated as the ratio of the normalized depth in males against the normalized depth in females. If the relative sequencing depth was around one, it suggested that there was no duplication of the GSD region in both females and males. If the relative sequencing depth was around two, it indicated a duplicated GSD locus in males. If it was about 0.5, it demonstrated that a duplicated GSD region might exist in females.

We further performed genotyping in the female population, the male population, two females and two males to fine map the GSD region. The cleaned genome re-sequencing reads of two populations and four individuals were aligned to the assembly with BWA (Li and Durbin 2010), respectively. Duplicated reads were removed using Picard in the GATK package (McKenna et al. 2010). After local realignment and base quality score recalibration, we retained those alignments of mapping score higher than 30. The variants and genotypes in each sample were detected using GATK. We observed that in the genetic map the sex-linked markers were consistently heterozygous in males but were homozygous in females. On the basis of this observation, among all re-sequenced sites we identified the sites which were heterozygous in all males but were homozygous in all females. These sites were considered to be sex-biased. Two sex-biased SNPs were selected to validate their segregation modes between two sexes (supplementary note 9). The functional effects of the associated SNPs on the corresponding gene were annotated by SnpEff (Cingolani et al. 2012) (supplementary note 10).

### Recombination rates of the GSD locus in males and females

The F1 cross was previously described in the construction of an integrated croaker genetic map (Xiao, Wang, et al. 2015). We first clustered all identified SNP markers of the mapping family by using JoinMap (Van Ooijen 2011). More markers that were grouped with the ones in LG9 of the integrated map were used to construct the female and male genetic map of LG9. Secondly, the SNP loci in this group were classified into three types: (aa × ab), (ab × aa) and (ab × ab). We used (aa × ab) and (ab × aa) markers to construct the female map and male map, respectively, using JoinMap (Van Ooijen 2011), respectively. After the orders of the (aa × ab) markers were fitted in the original female map, we combined the (ab × ab) markers together with (aa × ab) markers to re-construct the female map. Likewise, we constructed the male map using the markers of (ab × aa) and (ab × ab). The parameters of population type, mapping function, mapping algorithm and LOD threshold were used as our previous method (Xiao, Wang, et al. 2015). Thirdly, based on each map, we identified sex-linked markers by performing QTL analysis with the above parameters. The common markers shared by the male map and female maps were employed to compare the recombination rates around the GSD between males and females.

### FISH mapping

The chromosome slides prepared from head kidney cells of croaker were aged, denatured and dehydrated before hybridization. The DNA probes constructed from a BAC clone containing *dmrt1* (supplementary note 11) were labeled with biotin-11-dUTP using a Nick Translation System (Roche Diagnostics, Basel, Switzerland). The probes were added to the hybridization buffer, heated to be denatured together with sonicated Salmon sperm DNA, and then added to the denatured slides allowing hybridization in a moist chamber at 37 °C for at least 8 hours. After hybridization, the chromosome slides were washed serially with 50% formamide in 2 × SSC for 5 min at 42 °C, 2 × and 1 × SSC each for 5 min at room temperature. The biotinylated probes specifically attached to the slides were detected by avidin-Alexa Flour-488 (Invitrogen, CA, USA), and chromosomes were counter-stained with propidium iodide in anti-fade solution. The images of the chromosomes were observed and captured by an Olympus BX53 fluorescence microscope coupled with a micro CCD camera DP80, and analyzed with cellSens Standard 1.7 software.

### Differential expression analysis between males and females and between hormone-induced croaker sex reversals

Gonads of females, males and pseudo-males of two years old were observed under a microscope after processing by a standard histological protocol (supplementary note 12). In brief, gonads were dehydrated with a graded series of ethanol, equilibrated in xylene and embedded in paraffin. Slides with paraffin sections were deparaffinized, rehydrated and finally stained with hematoxylin and eosin (H&E). The stained sections were observed with a light microscope (Olympus, Tokyo, Japan).

The cleaned RNA-sequencing reads from gonads of mature females, mature males and pseudo-males were mapped to the assembly by bowtie2 (Langmead and Salzberg 2012). For each tissue, the read counts for each gene were calculated by HTSeq (Anders et al. 2015) and then normalized with edgeR (Robinson et al. 2010). FPKM (Fragments Per Kilobase of transcript per Million fragments) was applied to represent the normalized expression value. Three comparisons included females vs males, females vs pseudo-males, and males vs pseudo-males. For each comparison, we identified DEGs of fold change ≥ 4 and false discovery rate (FDR) ≤ 0.05. Since males and pseudo-males had the similar physiological characteristics of gonads, we used the DEGs in the comparison of males vs pseudo-males as false positive control and filtered them out of the DEGs in the comparison of females vs males and females vs pseudo-males. The retained DEGs were considered to be positive DEGs and used in the following analysis.

For each category of positive DEG, the GO enrichment was performed with hyper-geometry distribution test using GOATOOLS (Klopfenstein et al. 2018) software. We corrected the *p* value of each GO term for multiple testing with the Holm method (Holm 1979). The GO terms with corrected *p* ≤ 0.05 were considered to be statistically enriched in this group. We also investigated the pathways where the number of DEGs was significantly more than the number of all genes in the genome, assuming a hypergeometric distribution. The *p* value was adjusted with the Benjamin-Hochberg procedure (Benjamini and Hochberg 1995). Pathways with adjusted *p* value ≤ 0.05 were statistically enriched.

Transcriptome analysis identified 31 GSD genes having differential expression profiles among three types of gonads. The qRT-PCR analysis was performed on two GSD genes (*dmrt1* and *rnf*) to confirm the differential expressions (supplementary note 13). To identify co-expressed genes with *dmrt1*, a co-expression gene network for transcriptomes from males, females, and pseudo-males, was constructed using the R package WGCNA (supplementary note 14).

## Accession codes

The chromosome-level draft genome was submitted to the European Nucleotide Archive (ENA) with accession number of REGW00000000. All Illumina reads for genome assembly, re-sequencing, and RNA-seq were deposited at NCBI SRA database under accessions of PRJNA309029, PRJNA309060, and PRJNA368644. The Pacbio reads were available at SRA database under accessions of SRR7881518 and SRR7881519.

## Acknowledgements

This work was supported by grants from the National Natural Science Foundation of China (grant numbers U1205122, 31172397, 31402353, 31602207, and 31272653), Key projects of the Xiamen Southern Ocean Research Center (grant number 14GZY70NF34), and China Agriculture Research System (grant number CARS-47-G04).

## References

Aken BL, Achuthan P, Akanni W, Amode MR, Bernsdorff F, Bhai J, Billis K, Carvalho-Silva D, Cummins C, Clapham P, et al. 2017. Ensembl 2017. Nucleic Acids Res 45:D635–D642.

Anders S, Pyl PT, Huber W. 2015. HTSeq--a Python framework to work with high-throughput sequencing data. Bioinformatics 31:166–169.

Ao J, Mu Y, Xiang LX, Fan D, Feng M, Zhang S, Shi Q, Zhu LY, Li T, Ding Y, et al. 2015. Genome sequencing of the perciform fish Larimichthys crocea provides insights into molecular and genetic mechanisms of stress adaptation. PLoS Genet 11:e1005118.

Bao W, Kojima KK, Kohany O. 2015. Repbase Update, a database of repetitive elements in eukaryotic genomes. Mob DNA 6:11.

Benjamini Y, Hochberg Y. 1995. Controlling the False Discovery Rate: A Practical and Powerful Approach to Multiple Testing. Journal of the Royal Statistical Society. Series B (Methodological) 57:289–300.

Bergero R, Qiu S, Forrest A, Borthwick H, Charlesworth D. 2013. Expansion of the Pseudo-autosomal Region and Ongoing Recombination Suppression in the Silene latifolia Sex Chromosomes. Genetics 194:673.

Bogani D, Siggers P, Brixey R, Warr N, Beddow S, Edwards J, Williams D, Wilhelm D, Koopman P, Flavell RA, et al. 2009. Loss of Mitogen-Activated Protein Kinase Kinase Kinase 4 (MAP3K4) Reveals a Requirement for MAPK Signalling in Mouse Sex Determination. PLoS Biol 7:e1000196.

Brunner B, Hornung U, Shan Z, Nanda I, Kondo M, Zend-Ajusch E, Haaf T, Ropers H-H, Shima A, Schmid M, et al. 2001. Genomic Organization and Expression of the Doublesex-Related Gene Cluster in Vertebrates and Detection of Putative Regulatory Regions for DMRT1. Genomics 77:8–17.

Charlesworth B. 1991. The evolution of sex chromosomes. Science 251:1030–1033.

Chen S, Zhang G, Shao C, Huang Q, Liu G, Zhang P, Song W, An N, Chalopin D, Volff JN, et al. 2014. Whole-genome sequence of a flatfish provides insights into ZW sex chromosome evolution and adaptation to a benthic lifestyle. Nat Genet 46:253–260.

Chen Z WZ, Liu X, Jiang Y, Cai M. 2014. Area and physical length of metaphase chromosomes in large yellow croaker (Larimichthys crocea). J Fish China. 5:002.

Cingolani P, Platts A, Wang LL, Coon M, Nguyen T, Wang L, Land SJ, Lu X, Ruden DM. 2012. A program for annotating and predicting the effects of single nucleotide polymorphisms, SnpEff: SNPs in the genome of Drosophila melanogaster strain w1118; iso-2; iso-3. Fly 6:80–92.

Conesa A, Gotz S, Garcia-Gomez JM, Terol J, Talon M, Robles M. 2005. Blast2GO: a universal tool for annotation, visualization and analysis in functional genomics research. Bioinformatics 21:3674–3676.

Consortium TU. 2015. UniProt: a hub for protein information. Nucleic Acids Research 43:D204–D212.

Cox MP, Peterson DA, Biggs PJ. 2010. SolexaQA: At-a-glance quality assessment of Illumina second-generation sequencing data. BMC Bioinformatics 11:485.

Crews D, Bull JJ. 2009. Mode and tempo in environmental sex determination in vertebrates. Semin Cell Dev Biol 20:251–255.

Cui KH, Warnes GM, Jeffrey R, Matthews CD. 1994. Sex determination of preimplantation embryos by human testis-determining-gene amplification. The Lancet 343:79–82.

Edgar RC. 2004. MUSCLE: multiple sequence alignment with high accuracy and high throughput. Nucleic Acids Res 32:1792–1797.

Ferguson-Smith M. 2007. The evolution of sex chromosomes and sex determination in vertebrates and the key role of DMRT1. Sex Dev 1:2–11.

Fraser JA, Heitman J. 2005. Chromosomal sex-determining regions in animals, plants and fungi. Current Opinion in Genetics & Development 15:645–651.

Ghosh S, Chan CK. 2016. Analysis of RNA-Seq Data Using TopHat and Cufflinks. Methods Mol Biol 1374:339–361.

Guindon S, Lethiec F, Duroux P, Gascuel O. 2005. PHYML Online--a web server for fast maximum likelihood-based phylogenetic inference. Nucleic Acids Res 33:W557–559.

Hedges SB, Dudley J, Kumar S. 2006. TimeTree: a public knowledge-base of divergence times among organisms. Bioinformatics 22:2971–2972.

Hirakawa I, Miyagawa S, Mitsui N, Miyahara M, Onishi Y, Kagami Y, Kusano T, Takeuchi T, Ohta Y, Iguchi T. 2012. Developmental disorders and altered gene expression in the tropical clawed frog (Silurana tropicalis) exposed to 17α-ethinylestradiol. Journal of Applied Toxicology 33:1001–1010.

Holm S. 1979. A Simple Sequentially Rejective Multiple Test Procedure. Scandinavian Journal of Statistics 6:65–70.

Hong CS, Park BY, Saint-Jeannet JP. 2007. The function of Dmrt genes in vertebrate development: it is not just about sex. Dev Biol 310:1–9.

Howe K, Clark MD, Torroja CF, Torrance J, Berthelot C, Muffato M, Collins JE, Humphray S, McLaren K, Matthews L, et al. 2013. The zebrafish reference genome sequence and its relationship to the human genome. Nature 496:498–503.

Jordan CY, Charlesworth D. 2012. The potential for sexually antagonistic polymorphism in different genome regions. Evolution 66:505–516.

Kasahara M, Naruse K, Sasaki S, Nakatani Y, Qu W, Ahsan B, Yamada T, Nagayasu Y, Doi K, Kasai Y, et al. 2007. The medaka draft genome and insights into vertebrate genome evolution. Nature 447:714–719.

Kent WJ. 2002. BLAT—The BLAST-Like Alignment Tool. Genome Research 12:656–664.

Klopfenstein DV, Zhang L, Pedersen BS, Ramirez F, Warwick Vesztrocy A, Naldi A, Mungall CJ, Yunes JM, Botvinnik O, Weigel M, et al. 2018. GOATOOLS: A Python library for Gene Ontology analyses. Sci Rep 8:10872.

Kohno S, Bernhard MC, Katsu Y, Zhu J, Bryan TA, Doheny BM, Iguchi T, Guillette LJ. 2015. Estrogen Receptor 1 (ESR1; ERα), not ESR2 (ERβ), Modulates Estrogen-Induced Sex Reversal in the American Alligator, a Species With Temperature-Dependent Sex Determination. Endocrinology 156:1887–1899.

Komen H, Thorgaard GH. 2007. Androgenesis, gynogenesis and the production of clones in fishes: A review. Aquaculture 269:150–173.

Kondo M, Hornung U, Nanda I, Imai S, Sasaki T, Shimizu A, Asakawa S, Hori H, Schmid M, Shimizu N, et al. 2006. Genomic organization of the sex-determining and adjacent regions of the sex chromosomes of medaka. Genome Res 16:815–826.

Langfelder P, Horvath S. 2008. WGCNA: an R package for weighted correlation network analysis. BMC Bioinformatics 9:559.

Langmead B, Salzberg SL. 2012. Fast gapped-read alignment with Bowtie 2. Nat Methods 9:357–359.

Larkin MA, Blackshields G, Brown NP, Chenna R, McGettigan PA, McWilliam H, Valentin F, Wallace IM, Wilm A, Lopez R, et al. 2007. Clustal W and Clustal X version 2.0. Bioinformatics (Oxford, England) 23:2947–2948.

Li H, Durbin R. (r01820 co-authors). 2010. Fast and accurate long-read alignment with Burrows-Wheeler transform. Bioinformatics 26:589–595.

Li M, Sun Y, Zhao J, Shi H, Zeng S, Ye K, Jiang D, Zhou L, Sun L, Tao W. 2015. A Tandem Duplicate of Anti-Müllerian Hormone with a Missense SNP on the Y Chromosome Is Essential for Male Sex Determination in Nile Tilapia, Oreochromis niloticus. PLoS Genet 11:e1005678.

Liu Z, Liu S, Yao J, Bao L, Zhang J, Li Y, Jiang C, Sun L, Wang R, Zhang Y, et al. 2016. The channel catfish genome sequence provides insights into the evolution of scale formation in teleosts. Nat Commun 7:11757.

Livernois AM, Graves JA, Waters PD. 2012. The origin and evolution of vertebrate sex chromosomes and dosage compensation. Heredity (Edinb) 108:50–58.

Marçais G, Kingsford C. 2011. A fast, lock-free approach for efficient parallel counting of occurrences of k-mers. Bioinformatics (Oxford, England) 27:764–770.

Matsuda M, Nagahama Y, Shinomiya A, Sato T, Matsuda C, Kobayashi T, Morrey CE, Shibata N, Asakawa S, Shimizu N, et al. 2002. DMY is a Y-specific DM-domain gene required for male development in the medaka fish. Nature 417:559–563.

McKenna A, Hanna M, Banks E, Sivachenko A, Cibulskis K, Kernytsky A, Garimella K, Altshuler D, Gabriel S, Daly M. 2010. The Genome Analysis Toolkit: a MapReduce framework for analyzing next-generation DNA sequencing data. Genome Research 20:1297–1303.

Megosh HB, Cox DN, Campbell C, Lin H. 2006. The role of PIWI and the miRNA machinery in Drosophila germline determination. Curr Biol 16:1884–1894.

Moriya Y, Itoh M, Okuda S, Yoshizawa AC, Kanehisa M. 2007. KAAS: an automatic genome annotation and pathway reconstruction server. Nucleic Acids Res 35:W182–185.

Muller HJ. 1914. A gene for the fourth chromosome of Drosophila. Journal of Experimental Zoology 17:325–336.

Murata C, Kuroki Y, Imoto I, Tsukahara M, Ikejiri N, Kuroiwa A. 2015. Initiation of recombination suppression and PAR formation during the early stages of neo-sex chromosome differentiation in the Okinawa spiny rat, Tokudaia muenninki. BMC Evolutionary Biology 15:234.

Nanda I, Kondo M, Hornung U, Asakawa S, Winkler C, Shimizu A, Shan Z, Haaf T, Shimizu N, Shima A. 2002. A duplicated copy of DMRT1 in the sex-determining region of the Y chromosome of the medaka, Oryzias latipes. Proceedings of the National Academy of Sciences 99:11778–11783.

Narita S, Kageyama D, Nomura M, Fukatsu T. 2007. Unexpected mechanism of symbiont-induced reversal of insect sex: feminizing Wolbachia continuously acts on the butterfly Eurema hecabe during larval development. Appl Environ Microbiol 73:4332–4341.

Nef S, Verma-Kurvari S, Merenmies J, Vassalli J-D, Efstratiadis A, Accili D, Parada LF. 2003. Testis determination requires insulin receptor family function in mice. Nature 426:291.

Ning Y, Liu X, Wang ZY, Guo W, Li Y, Xie F. 2007. A genetic map of large yellow croaker Pseudosciaena crocea. Aquaculture 264:16–26.

Otani A, Nakajima T, Okumura T, Fujii S, Tomooka Y. 2017. Sex Reversal and Analyses of Possible Involvement of Sex Steroids in Scallop Gonadal Development in Newly Established Organ-Culture Systems. Zoological Science 34:86–92.

Parra G, Bradnam K, Korf I. 2007. CEGMA: a pipeline to accurately annotate core genes in eukaryotic genomes. Bioinformatics 23:1061–1067.

Peichel CL, Ross JA, Matson CK, Dickson M, Grimwood J, Schmutz J, Myers RM, Mori S, Schluter D, Kingsley DM. 2004. The master sex-determination locus in threespine sticklebacks is on a nascent Y chromosome. Curr Biol 14:1416–1424.

Phillips RB, DeKoning JJ, Ventura AB, Nichols KM, Drew RE, Chaves LD, Reed KM, Felip A, Thorgaard GH. 2009. Recombination is suppressed over a large region of the rainbow trout Y chromosome. Animal Genetics 40:925–932.

Piferrer F. 2001. Endocrine sex control strategies for the feminization of teleost fish. Aquaculture 197:229–281.

Pradhan A, Olsson PE. 2015. Zebrafish sexual behavior: role of sex steroid hormones and prostaglandins. Behav Brain Funct 11:23.

Price AL, Jones NC, Pevzner PA. 2005. De novo identification of repeat families in large genomes. Bioinformatics 21 Suppl 1:i351–358.

Purcell S, Neale B, Todd-Brown K, Thomas L, Ferreira MA, Bender D, Maller J, Sklar P, de Bakker PI, Daly MJ, et al. 2007. PLINK: a tool set for whole-genome association and population-based linkage analyses. Am J Hum Genet 81:559–575.

Reisser CM, Fasel D, Hurlimann E, Dukic M, Haag-Liautard C, Thuillier V, Galimov Y, Haag CR. 2017. Transition from Environmental to Partial Genetic Sex Determination in Daphnia through the Evolution of a Female-Determining Incipient W Chromosome. Mol Biol Evol 34:575–588.

Rice WR. 1987. The accumulation of sexually antagonistic genes as a selective agent promoting the evolution of reduced recombination between primitive sex chromosomes. Evolution 41:911–914.

Robinson MD, McCarthy DJ, Smyth GK. 2010. edgeR: a Bioconductor package for differential expression analysis of digital gene expression data. Bioinformatics 26:139–140.

Schmidt EE, Schibler U. 1997. Developmental testis-specific regulation of mRNA levels and mRNA translational efficiencies for TATA-binding protein mRNA isoforms. Developmental biology 184:138–149.

Schreiber F, Patricio M, Muffato M, Pignatelli M, Bateman A. 2014. TreeFam v9: a new website, more species and orthology-on-the-fly. Nucleic Acids Res 42:D922–925.

Singer A, Perlman H, Yan Y, Walker C, Corley-Smith G, Brandhorst B, Postlethwait J. 2002. Sex-specific recombination rates in zebrafish (Danio rerio). Genetics 160:649–657.

Smith CA, Katz M, Sinclair AH. 2003. DMRT1 is upregulated in the gonads during female-to-male sex reversal in ZW chicken embryos. Biol Reprod 68:560–570.

Smith CA, Roeszler KN, Ohnesorg T, Cummins DM, Farlie PG, Doran TJ, Sinclair AH. 2009. The avian Z-linked gene DMRT1 is required for male sex determination in the chicken. Nature 461:267–271.

Spigler RB, Lewers KS, Johnson AL, Ashman TL. 2010. Comparative mapping reveals autosomal origin of sex chromosome in octoploid Fragaria virginiana. J Hered 101 Suppl 1:S107–117.

Stelkens RB, Wedekind C. 2010. Environmental sex reversal, Trojan sex genes, and sex ratio adjustment: conditions and population consequences. Mol Ecol 19:627–646.

Suyama M, Torrents D, Bork P. 2006. PAL2NAL: robust conversion of protein sequence alignments into the corresponding codon alignments. Nucleic Acids Res 34:W609–612.

Takehana Y, Matsuda M, Myosho T, Suster ML, Kawakami K, Shin IT, Kohara Y, Kuroki Y, Toyoda A, Fujiyama A, et al. 2014. Co-option of Sox3 as the male-determining factor on the Y chromosome in the fish Oryzias dancena. Nat Commun 5:4157.

Talavera G, Castresana J. 2007. Improvement of phylogenies after removing divergent and ambiguously aligned blocks from protein sequence alignments. Syst Biol 56:564–577.

Tennessen JA, Govindarajulu R, Liston A, Ashman TL. 2016. Homomorphic ZW chromosomes in a wild strawberry show distinctive recombination heterogeneity but a small sex-determining region. New Phytologist 211:1412–1423.

Toshima J, Ohashi K, Okano I, Nunoue K, Kishioka M, Kuma K, Miyata T, Hirai M, Baba T, Mizuno K. 1995. Identification and characterization of a novel protein kinase, TESK1, specifically expressed in testicular germ cells. J Biol Chem 270:31331–31337.

Trapnell C, Pachter L, Salzberg SL. 2009. TopHat: discovering splice junctions with RNA-Seq. Bioinformatics 25:1105–1111.

Vallender EJ, Lahn BT. 2006. Multiple independent origins of sex chromosomes in amniotes. Proc Natl Acad Sci U S A 103:18031–18032.

Van Ooijen J, Kyazma B. 2009. MapQTL 6. Software for the mapping of quantitative trait loci in experimental populations of diploid species. Kyazma BV: Wageningen, Netherlands.

Van Ooijen JW. 2011. Multipoint maximum likelihood mapping in a full-sib family of an outbreeding species. Genetics Research 93:343–349.

Wang Y, Lu Y, Zhang Y, Ning Z, Li Y, Zhao Q, Lu H, Huang R, Xia X, Feng Q, et al. 2015. The draft genome of the grass carp (Ctenopharyngodon idellus) provides insights into its evolution and vegetarian adaptation. Nat Genet 47:625.

Wang Y, Tang H, DeBarry JD, Tan X, Li J, Wang X, Lee T-h, Jin H, Marler B, Guo H. 2012. MCScanX: a toolkit for detection and evolutionary analysis of gene synteny and collinearity. Nucleic Acids Research 40:e49–e49.

Wu C, Zhang D, Kan M, Lv Z, Zhu A, Su Y, Zhou D, Zhang J, Zhang Z, Xu M, et al. 2014. The draft genome of the large yellow croaker reveals well-developed innate immunity. Nature communications 5:5227.

Xiao CL, Chen Y, Xie SQ. 2017. MECAT: fast mapping, error correction, and de novo assembly for single-molecule sequencing reads. 14:1072–1074.

Xiao S, Han Z, Wang P, Han F, Liu Y, Li J, Wang ZY. 2015. Functional marker detection and analysis on a comprehensive transcriptome of large yellow croaker by next generation sequencing. PLoS One 10:e0124432.

Xiao S, Wang P, Zhang Y, Fang L, Liu Y, Li J-T, Wang Z-Y. 2015. Gene map of large yellow croaker (Larimichthys crocea) provides insights into teleost genome evolution and conserved regions associated with growth. Scientific Reports 5:18661.

Yang Z. 2007. PAML 4: Phylogenetic Analysis by Maximum Likelihood. Mol Biol Evol 24:1586–1591.

