## Supplementary text for "Proto-sex locus in large yellow croaker provides insights into early evolution of the sex chromosome"

Table of Contents

1. Sample preparation for genomic sequencing	4

2. Whole genome sequencing and re-sequencing	5

3. Production of pseudo-males of croaker	5

4. Sequencing of transcriptome from gonads of males, females and pseudo-males	5

5. Estimation of the genome size	6

6. Genome assembly and anchoring scaffolds to the linkage map	6

7. Gene prediction and annotation	7

8. PCR amplification of randomly selected genes to assess the quality of croaker genome assembly	8

9. Validating the segregation modes of two sex-linked SNP markers	8

10. Annotation of SNP functions on *dmrt1* in croaker	9

11. Screening a sex-related BAC clone for FISH experiments	10

12. Morphological examination of gonads from females, males and pseudo-males	10

13. Validation of two DEGs expression using qRT-PCR	11

References	12

Supplementary Figures	14

FIG. S1. Estimating genome size with K-mer distribution	14

FIG. S2. The chromosome-level assembly of the croaker	15

FIG. S3. Distributions of insert sizes of sequencing reads of different libraries on the genome assembly	16

FIG. S4. Comparison of gene structure of nine teleost species	17

FIG. S5. Amplification of 24 novel genes	18

FIG. S6. Orthologous profiling among croaker, medaka, stickleback, European sea bass and tetraodon	19

FIG. S7. Correlation between intergenic length and genome size and between the proportions of different types of repeats and genome size	20

FIG. S8. Sanger sequencing of two SNPs segregated perfectly with females and males.	22

FIG. S9. K-mer distribution of four individuals	23

FIG. S10. The phylogenetic tree of *dmrt* family in vertebrates	24

FIG. S11. Profiling of *dmrt1* expression across tissues	25

FIG. S12. qRT-PCR validation of two GSD DEGs in males, females and pseudo-males	26

FIG. S13. Profiling *dmrt1* expression across developmental stages	27

FIG. S14. Sex-biased markers in the transcription factor binding sites of *dmrt1*.	27

FIG. S15. Overlapping of three groups of DEGs.	28

FIG. S16. Significantly co-expressed genes with *dmrt1* identified by WGCNA	28

Supplementary Tables	29

Table S1. Summary of library construction and sequencing	29

Table S2. Summary of croaker genome assembly	29

Table S3. The quality metrics of previous two assemblies and our assembly	29

Table S4. Mapping ratio of Illumina genomic sequencing reads to our croaker assembly	30

Table S5. Mapping ratio of re-sequencing reads to our croaker assembly	30

Table S6. Mapping ratio of CEGMA proteins to our croaker assembly	30

Table S7. Summary of croaker genome components	30

Table S8. The statistics of gene structures of croaker and other teleosts	31

Table S9. Redundant genes in the previous assemblies	32

Table S10. Annotations of croaker genes against known protein databases	33

Table S11. Primers for amplification of twenty-four randomly selected novel genes	33

Table S12. Assignment ratio of teleost genes to TreeFam families	35

Table S14. Repeat content in croaker genome	37

Table S15. Interspersed repeat components of eight teleost genomes	38

Table S16. Six sex-linked markers in the integrated chr9 of croaker	39

Table S17. 617 sites with polymorphism segregation in two sexes	40

Table S18. The functions of 79 genes in croaker GSD locus	40

Table S19. Primers used to validate two sex-biased markers.	40

Table S20. Estimated genome sizes of four individuals	40

Table S21. The sex-biased SNPs and Ks of genes in the GSD locus	41

Table S22. The predicted biological functions of the sex-biased SNPs and InDels around *dmrt1*	42

Table S23. Cleaned RNA-seq reads of male, pseudo-male and female gonads of croaker	43

Table S24. Expression levels of 31 DEGs in the GSD among males, females and pseudo-males	44

Table S25. Primers of two DEGs for qRT-PCR analysis.	45

Table S26. The functions of 66 genes significantly co-expressed with *dmrt1*.	45

Table S27. GO and KEGG pathway enrichment for hormone-induce DEGs	45

Table S28. GO and KEGG pathway enrichment for genetic-determining specific DEGs	45

Table S29. The GO and KEGG pathways by common DEGs shared in both GSD and hormone-induced ESR	45

**Supplementary Note**

#### 1. Sample preparation for genomic sequencing

To produce a high-quality genome assembly, one individual, a gynogen of second generation of meiotic gynogenesis, was collected from the Fishery Technical Extension Station of Ningde City, Fujian province, China. The first artificial meiotic gynogenesis was carried out in spring, 2001. Briefly, mature fish were selected from a culture population. Oviposition was induced by injecting a GnRH analogue (Ningbo Hormone Plant, Zhejiang, China) with a dosage of 3 μg per kilogram of body weight. Eggs and semen were collected approximately 36 h after the injection. We then followed Wang *et al*.’s strategy to induce gynogenesis. The semen was treated with ultraviolet light (wavelength: 254 nm). Then the eggs were inseminated with irradiated sperm. After 3 minutes post-insemination, the extrusion of the second polar body was blocked by cold shock at 3 °C for 10 minutes. In spring, 2003, the mature gynogens of the first generation were selected and induced to oviposit using the GnRH analogue eggs, and eggs were collected. Semen of mature males from the other culture population was inactivated with ultraviolet light. Then the eggs were inseminated with the treated sperm and the extrusion of the second polar body was blocked following the above-mentioned method. In spring, 2007, the fins of 48 gynogens of the second generation were collected and genomic DNA was extracted from the fins. We used 15 highly polymorphic microsatellite markers of croaker to investigate the homozygosity of these gynogens. The homozygosity of 48 gynogens ranged from 0.600 to 0.933 . The individual with the highest homozygosity was selected for whole genome sequencing.

In addition, DNA from dorsal fins of fifty males were collected and pooled together. This male pool was re-sequenced for further analysis. Likewise, the DNA of a female pool comprising fifty females was also prepared for genome re-sequencing. Finally, we also extracted genomic DNAs from additional two randomly sampled females (female 1 and female 2) and two randomly sampled males (male 1 and male 2) from a culture population for re-sequencing, respectively.

#### 2. Whole genome sequencing and re-sequencing

For *de novo* genome sequencing, we constructed five sequencing libraries including short paired-end libraries with insert sizes of 200 bp and 600 bp, and mate-paired libraries spanning genomic distances of 1 kb, 6 kb and 8 kb. The five libraries were sequenced with 100 PE mode on Illumina platform following standard manufacturer’s protocols (supplementary table S1). The genomic DNA molecules were also sequenced using PacBio Sequel platform. Two 20 kb libraries were constructed and then sequenced on the Sequel platform.

We constructed paired-end re-sequencing libraries with the insert size of 300 bp for the male population, female population, and four randomly sampled individuals, respectively. The six re-sequencing libraries were also sequenced on Illumina platform.

#### 3. Production of pseudo-males of croaker

Artificial gynogenetic diploids were produced and screened according to the method described by our previous studies . About 5,000 gynogenetic larvae were subject to hormone treatment to produce pseudo-males during the liable period of sex reversal. The individuals were fed diet with 17α-methyltestosterone (5 mg/kg body weight) from 30 days after hatching (dph) to 120 dph and then switched to normal diet. The gender of each mature individual was identified by examining the extrusion from the gonad or dissection.

#### 4. Sequencing of transcriptome from gonads of males, females and pseudo-males

Total RNA was extracted from gonad tissues of five mature females, three males and three pseudo-males with Total RNA kit II (OmegaBioteck, Georgia, USA). The concentration of RNA was measured using Qubit RNA Assay Kit on Qubit 2.0 Flurometer (Life Technologies, CA, USA) and the integrity were assessed using 2100 Bioanalyzer (Agilent Technologies, CA, USA). The RNA-seq library with insert length of 300 bp for each sample was constructed following the standard protocol. The libraries were sequenced on the Illumina platform with 125 bp pair-end mode (Illumina, CA, USA).

#### 5. Estimation of the genome size

The cleaned paired-end reads of the two paired-end libraries (200 bp and 600 bp) from the sequenced gynogen were used to estimate genome size of croaker. We estimated the frequency of 17 bp K-mer using Jellyfish . Following Star *et al.*’s method, the croaker genome size was estimated by dividing the total amount of sequenced bases with sequencing depth.

We also estimated the genome sizes of the sequenced two females and two males using each short-insert paired-end library with the above-mentioned method.

#### 6. Genome assembly and anchoring scaffolds to the linkage map

After assembling the cleaned pair-end reads to contigs using SOAPdenovo with default parameters, reads from mate-pairs of different insert sizes were step-by-step added for scaffolding contigs using SSPACE , Opera and SOPRA . The assembled croaker transcriptome from multiple tissues was used to further scaffold the assembly using L_RNA_scaffolder , resulting in the improved genome assembly. We used the paired-end information from the short paired-end reads to fill the gaps between the scaffolds with Gapcloser and Gapfiller . Finally, the error-corrected Pacbio reads were utilized to close the gaps in the assembly and improve the contiguity using LR_Gapcloser (http://www.fishbrowser.org/software/LR_Gapcloser/).

A genetic map consisting of 3,448 SNP markers was used to anchor the scaffolds into a chromosome-level assembly. The flanking sequences of markers were aligned to the scaffolds using BLAT . The flanking sequence of a marker aligned at least 70% of its bases to a specific scaffold were kept. For those markers, which could be aligned to multiple loci, only the best alignment (with longest aligned bases) was retained for further analysis. If one marker had more than two equally best-aligned regions, the marker and corresponding alignment regions were discarded. When a scaffold was anchored to multiple linkage groups by different markers, the localization with more markers was retained.

#### 7. Gene prediction and annotation

Three approaches, including *de novo* gene prediction, homology-based prediction and RNA-seq models were adopted to annotate genes. Firstly, Fgenesh, a *de novo* prediction package , was used to predict genes on repeat-masked genome sequence. Secondly, teleost proteins in Ensembl database were aligned to croaker genome using BLAT . We retained alignments where over 70% of amino acids of a specific protein can be aligned. The proteins were re-aligned to these genome fragments by GeneWise for accurately spliced align­ments. Thirdly, RNA-seq reads from multiple tissues (embryo, liver, spleen, brain, kidney and head kidney) were mapped to genomic sequences using Tophat and then Cufflinks was used to assemble transcripts. All three sets of gene models were merged to form a comprehensive consensus gene set using cuffmerge . In case of consensus genes with several alternative splicing transcripts, the longest transcripts were chosen to represent those genes. The protein sequences of the representative transcripts were predicted using Transdecoder (https://transdecoder.github.io/).

To compare the gene structures of croaker with those in other vertebrates, including zebrafish, Atlantic cod, stickleback, medaka, European sea bass, takifugu, tetraodon, chicken, and human, we downloaded gene sets of these species from Ensembl and European sea bass genome database (http://seabass.mpipz.mpg.de). We used the longest protein-coding transcript to represent each gene in these species to compare the coding length, intergenic length, exon number per gene, exon size, and intron size among them.

#### 8. PCR amplification of randomly selected genes to assess the quality of croaker genome assembly

We aligned all predicted genes in our assembly to previous two assemblies using Blastn with e-value of 10^-5^. If predicted genes were not aligned, then they were considered as novel genes. To examine the quality of the croaker assembly, we used 24 randomly selected novel genes for PCR validation. The primers for amplifications were listed in supplementary table S11. PCR amplifications were performed in 20 μL solutions, including 1 μL (40 ng) of genome DNA, 2 μL of 10 × PCR buffer, 1 μL of 15 mmol/L MgCl_2_, 1.6 μL of 10 mmol/L dNTPs, 1 μL of 10 mmol/L primers and 0.2 μL of 5 U Hotstar Taq (Qiagen, Hilden, Germany). The reaction was performed on a PCR instrument (Eppendorf, Hamburg, Germany) with the following protocol: denaturation for 5 min at 94 °C; 45 cycles of 30 s at 94 °C, 30 s at annealing temperature, 30 s min at 72 °C, and a final extension at 72 °C for 10 min.

#### 9. Validating the segregation modes of two sex-linked SNP markers

To validate the segregation mode of sex-linked SNPs in more fish, two SNPs perfectly segregated with sex were selected and genotyped in another croaker population including 15 females and 15 males. The primers were shown in supplementary table S19**.** Genomic DNA was extracted from dorsal fins of the males and females following the method in supplementary note 1. PCR amplification was performed in a 5 μL solution containing 20 ng of genomic DNA, 0.5 U Hotstar Taq (Qiagen, Hilden, Germany), 0.5 μL 10 × PCR buffer, 0.1 μL dNTPs and 0.5 pmol of each primer. The PCR were carried out using the following steps: 4 min of denaturation at 94 ℃, 45 cycles of 20 sec at 94℃, 30 s at 56 ℃ and 1 min at 72 ℃ and a final extension at 72 ℃ for 3 min. The amplification products were sequenced using ABI Prism 3730 DNA Analyzer. The high quality sequences were aligned to the reference genome to examine the genotypes the SNPs.

#### 10. Annotation of SNP functions on *dmrt1* in croaker

We identified 34 SNPs segregated in the investigated croaker population in the *dmrt1* gene plus 3 kb upstream and 3 kb downstream (supplementary table S22). There were no variants found on the coding sequences of *dmrt1*. The 19 intronic SNPs were examined to see whether there were splicing site variants using SnpEff . As we know, SNPs in the miRNA binding site might interrupt the miRNA-mRNA interaction . We then retrieved the 3’UTR sequence of *dmrt1* and found that six SNPs located on the 3’UTR of this gene. In the 199 known croaker miRNAs , we predicted binding sites of these miRNA on the mutant and wild-type sequence of the 3’UTR using RNAhybrid with the default parameters. We compared the binding sites between wild-type and mutant 3’UTR. For the SNPs on the 5’UTR and upstream and downstream region of *dmrt1*, we predicted the possible transcriptional factor binding sites with an online prediction tools, ConSite , with conservation cutoff of 74%, window size of 20 bp, and 80% TF score threshold.

#### 11. Screening a sex-related BAC clone for FISH experiments

To identify and compare croaker sex chromosomes between males and females, we performed FISH experiments on head kidney cells from males and females. We constructed a three-dimension bacterial artificial chromosome (BAC) pool of croaker genome, containing 41,472 individual clones and corresponding to 8.1 × haploid genome coverage. We scanned BAC clones containing the sex-related genes in the GSD locus from the BAC library. The GSD locus comprised 79 genes, including *dmrt1* (supplementary table S18). To identify the BAC clone containing *dmrt1*, we designed a pair of PCR primers (F: 5´-CCAGCGAGACACCATACACC-3´ and R: 5´-GGCCATGAAACAGACGAAAG-3´) targeting *dmrt1* and its flanking genomic regions. A candidate positive clone was selected from the BAC library using this primer pair. Then the DNA probes constructed from the BAC clone were labeled with biotin-11-dUTP and hybridized with the chromosome slides prepared from head kidney cells of croaker, described in the Materials and Methods.

#### 12. Morphological examination of gonads from females, males and pseudo-males

The gonads of females, males and pseudo-males, stored in Bouin’s fixative, were compared to examine the morphological difference among them. The tissues were equilibrated in xylene and embedded in paraffin. After three hours of embedding in room temperature, sections of 6 μm were sliced with rotary microtome (KEDEE, Zhejiang, China) and mounted in glass slides. After proper rehydration treatment, the sections on the glass slides were then stained with H&E solutions. After three hours of incubation in room temperature, the stained histological slides were observed with a light microscope (Olympus, Tokyo, Japan), and the images were taken using CellSens standard imaging software (Olympus, Tokyo, Japan).

#### 13. Validation of two DEGs expression using qRT-PCR

We study the temporal and spatial expression patterns of *dmrt1*. Croakers were obtained from the Fishery Technology Extension Station of Ningde, Fujian, China. Larvae were collected at 6, 13, 20, 27, 34, 41, 49 and 55 dph. From 55 dph onwards, the gonads of juvenile (n > 10) were collected every two weeks up to 123 dph. The gonads of males and females at 8 months, 16 months and 2 years were also collected. Meanwhile, 11 tissues (eye, brain, liver, heart, kidney, gonad, head kidney, muscle, spleen, stomach and intestine) of both genders were dissected from healthy and adult croaker at age of 2 years (female 508 ± 88 g, male 450 ± 75 g) and were frozen in RNAlater immediately for followed RNA extraction and qRT-PCR. Total RNA was extracted as described in the previous section of RNA sequencing library preparation. The concentration of each RNA sample was adjusted to 1 μg/μL with nuclease-free water, and 2 μg of total RNA was reverse transcribed in a 20 μL reaction system using the TIANSsript RT Kit (TIANGEN, Beijing, China). In addition, we validated the expression patterns of two DEG (*dmrt1* and *rnf*) among gonads from males, females and pseudo-males.

For qRT-PCR experiment, the *β*-actin gene of croaker was used as an internal control. The qRT-PCR mixture had a total volume of 20 μL, including 10 μL 2 × TransStart Top Green qRT-PCR SuperMix (TRANsGen, Beijing, China), 0.4 μL of each primer (10 μM),1μL of cDNA, and 8.4 μL of RNase-free water. The reactions were performed on ABI 7300 Real Time System. The qRT-PCR program started with 30 s at 94 °C, followed by 40 cycles of 94 °C for 5 s and 60 °C for 31 seconds. The expression results were normalized to the expression level of the *β*-actin gene. A negative control (nuclease free water) was included in the experiment to detect whether there existed contamination. The specificity of qRT-PCR primers was confirmed by melting curve and sequencing of qRT-PCR products (supplementary table S25). The expressed quantitative variation was calculated as 2^-ΔΔCt^ .

To study the temporal expression pattern, the expression level in male’s gonad at 41 dph was set as a reference, because this was the time when *dmrt1* gene expression could be detected in male gonads. To study the spatial expression pattern of *dmrt1*, the expression level of this gene in female’s gonad was set as a reference. For *rnf*, the lowest expression level was set as a reference.

**14. Network analysis to identify co-expressed genes with *dmrt1* by WGCNA**

For transcriptomes from males, females, and pseudo-males, an unsigned co-expression gene network was constructed using the R package WGCNA with the parameters of ‘sft = 6, minimum module size = 30 and cutting height = 0.25’. Genes having similar expression profiles are grouped into a module. If certain genes have similar expression changes associated with the gender of samples, these genes are believed to be functionally related and can be defined as a module. The co-expressed genes, the weights of edges between which and *dmrt1* were over 0.7, were selected, and the co-expression interactions were constructed via Cytoscape .

### References

Aken BL, Achuthan P, Akanni W, Amode MR, Bernsdorff F, Bhai J, Billis K, Carvalho-Silva D, Cummins C, Clapham P, et al. 2017. Ensembl 2017. Nucleic Acids Res 45:D635-D642.

Ao J, Mu Y, Xiang LX, Fan D, Feng M, Zhang S, Shi Q, Zhu LY, Li T, Ding Y, et al. 2015. Genome sequencing of the perciform fish Larimichthys crocea provides insights into molecular and genetic mechanisms of stress adaptation. PLoS Genet 11:e1005118.

Birney E, Clamp M, Durbin R. 2004. GeneWise and Genomewise. Genome Res 14:988-995.

Boetzer M, Henkel CV, Jansen HJ, Butler D, Pirovano W. 2011. Scaffolding pre-assembled contigs using SSPACE. Bioinformatics 27:578-579.

Cingolani P, Platts A, Wang LL, Coon M, Nguyen T, Wang L, Land SJ, Lu X, Ruden DM. 2012. A program for annotating and predicting the effects of single nucleotide polymorphisms, SnpEff: SNPs in the genome of Drosophila melanogaster strain w1118; iso-2; iso-3. Fly 6:80-92.

Dayarian A, Michael TP, Sengupta AM. 2010. SOPRA: Scaffolding algorithm for paired reads via statistical optimization. BMC Bioinformatics 11:345.

Gao S, Sung WK, Nagarajan N. 2011. Opera: reconstructing optimal genomic scaffolds with high-throughput paired-end sequences. J Comput Biol 18:1681-1691.

Ghosh S, Chan CK. 2016. Analysis of RNA-Seq Data Using TopHat and Cufflinks. Methods Mol Biol 1374:339-361.

Huang Y, Cheng JH, Luo FN, Pan H, Sun XJ, Diao LY, Qin XJ. 2016. Genome-wide identification and characterization of microRNA genes and their targets in large yellow croaker (Larimichthys crocea). Gene 576:261-267.

Kent WJ. 2002. BLAT—The BLAST-Like Alignment Tool. Genome Research 12:656-664.

Kruger J, Rehmsmeier M. 2006. RNAhybrid: microRNA target prediction easy, fast and flexible. Nucleic Acids Res 34:W451-454.

Langfelder P, Horvath S. 2008. WGCNA: an R package for weighted correlation network analysis. BMC Bioinformatics 9:559.

Li Y, Cai M, Wang Z, Guo W, Liu X, Wang X, Ning Y. 2008. Microsatellite–centromere mapping in large yellow croaker (Pseudosciaena crocea) using gynogenetic diploid families. Marine Biotechnology 10:83-90.

Luo R, Liu B, Xie Y, Li Z, Huang W, Yuan J, He G, Chen Y, Pan Q, Liu Y, et al. 2012. SOAPdenovo2: an empirically improved memory-efficient short-read de novo assembler. Gigascience 1:18.

Marcais G, Kingsford C. 2011. A fast, lock-free approach for efficient parallel counting of occurrences of k-mers. Bioinformatics 27:764-770.

Nadalin F, Vezzi F, Policriti A. 2012. GapFiller: a de novo assembly approach to fill the gap within paired reads. BMC Bioinformatics 13 Suppl 14:S8.

Salamov AA, Solovyev VV. 2000. Ab initio gene finding in Drosophila genomic DNA. Genome Res 10:516-522.

Sandelin A, Wasserman WW, Lenhard B. 2004. ConSite: web-based prediction of regulatory elements using cross-species comparison. Nucleic Acids Res 32:W249-252.

Su G, Morris JH, Demchak B, Bader GD. 2014. Biological network exploration with Cytoscape 3. Curr Protoc Bioinformatics 47:8.13.11-24.

Trapnell C, Pachter L, Salzberg SL. 2009. TopHat: discovering splice junctions with RNA-Seq. Bioinformatics 25:1105-1111.

VanGuilder HD, Vrana KE, Freeman WM. 2008. Twenty-five years of quantitative PCR for gene expression analysis. Biotechniques 44:619.

Wang X, Wang Z, Liu X, Xie F, Liu J. 2006. Microsatellite marker analysis of gynogenesis by artificial induction in Pseudosciaena crocea. Yi chuan = Hereditas 28:831-837.

Wei Xue J-TL, Ya-Ping Zhu, Guang-Yuan Hou, Xiang-Fei Kong, You-Yi Kuangand Xiao-Wen Sun. 2013. L_RNA_scaffolder: scaffolding genomes with transcripts. BMC Genomics 14:604-617.

Wu C, Zhang D, Kan M, Lv Z, Zhu A, Su Y, Zhou D, Zhang J, Zhang Z, Xu M, et al. 2014. The draft genome of the large yellow croaker reveals well-developed innate immunity. Nature communications 5:5227.

Xiao S, Han Z, Wang P, Han F, Liu Y, Li J, Wang ZY. 2015. Functional marker detection and analysis on a comprehensive transcriptome of large yellow croaker by next generation sequencing. PLoS One 10:e0124432.

Xiao S, Wang P, Zhang Y, Fang L, Liu Y, Li JT, Wang ZY. 2015. Gene map of large yellow croaker (Larimichthys crocea) provides insights into teleost genome evolution and conserved regions associated with growth. Sci Rep 5:18661.

Ye X, Wang Z, Liu X, Cai M, Yao C. 2010. Analysis of genetic homozygosity and diversity of two successive generation meio-gynogenetic population in pseudosciaena crocea using microsatellite markers. Acta Hydrobiologica Sinica 34:144-151.

Zhu YP, Xue W, Wang JT, Wan YM, Wang SL, Xu P, Zhang Y, Li JT, Sun XW. 2012. Identification of common carp (Cyprinus carpio) microRNAs and microRNA-related SNPs. BMC Genomics 13:413.

### Supplementary Figures

#### FIG. S1. Estimating genome size with K-mer distribution

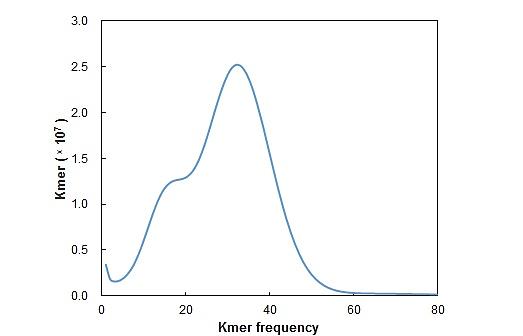

Distribution profile of unique K-mer counts in the cleaned sequencing reads of the two paired-end libraries. The K-mer size was 17.

#### FIG. S2. The chromosome-level assembly of the croaker

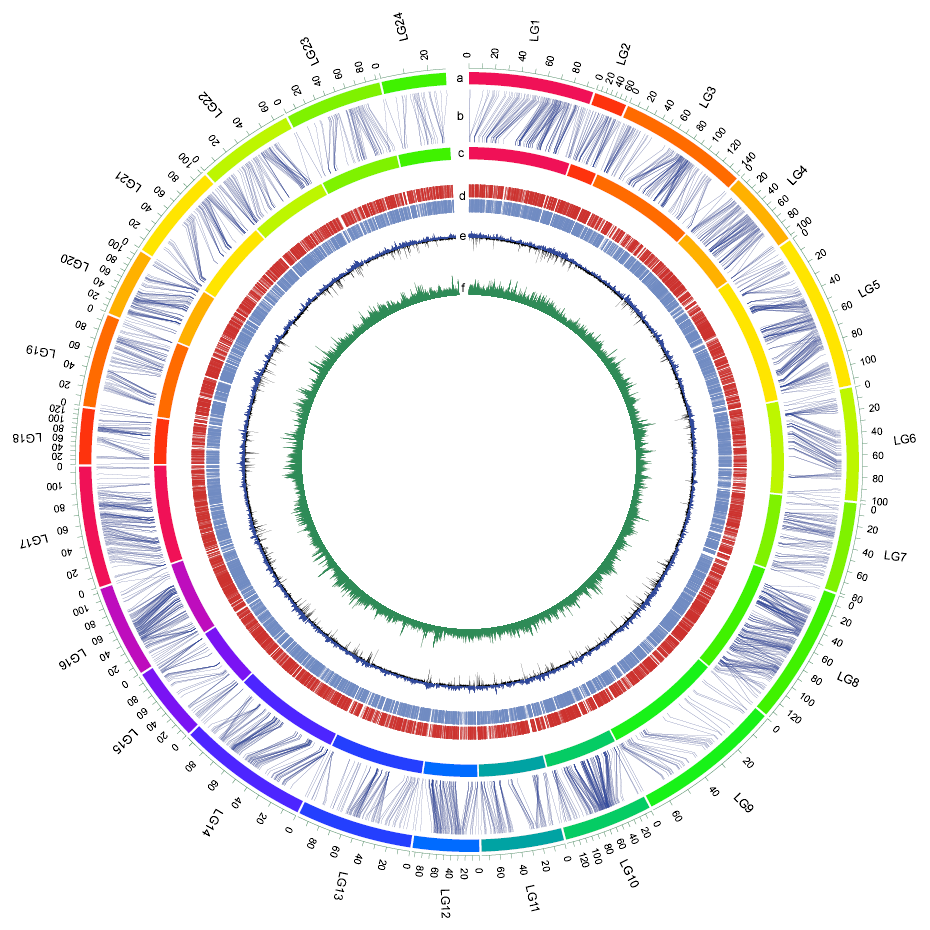

(a) The genetic linkage map. (b) Anchors between the genetic markers and the assembled scaffolds. (c) Assembled chromosomes. (d) Gene distribution on each chromosome; red lines indicate genes on the plus strand, and blue lines indicate genes on the minus strand. (e) GC content within a 10-kb sliding window; blue indicates GC content that is higher than average, and black indicates GC content that is lower than average. (f) Repeat content within a 10-kb sliding window.

#### FIG. S3. Distributions of insert sizes of sequencing reads of different libraries on the genome assembly

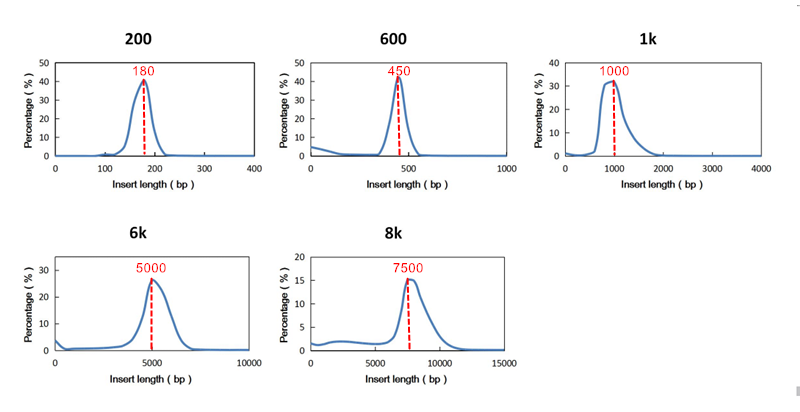

We aligned two ends of each pair from different paired-end/mate-pair libraries using BWA (settings: both reads in a pair should be uniquely mapped). If two ends of one pair were aligned to one genomic sequence, we calculated the distance between them.

#### FIG. S4. Comparison of gene structure of nine teleost species

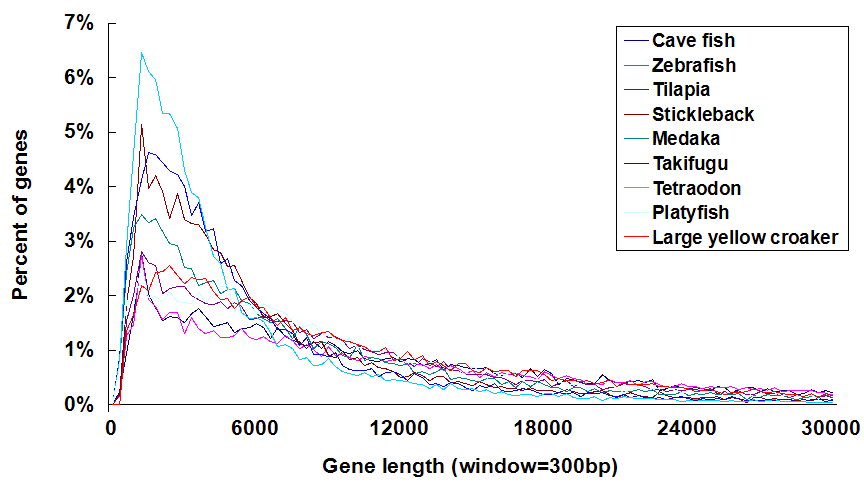

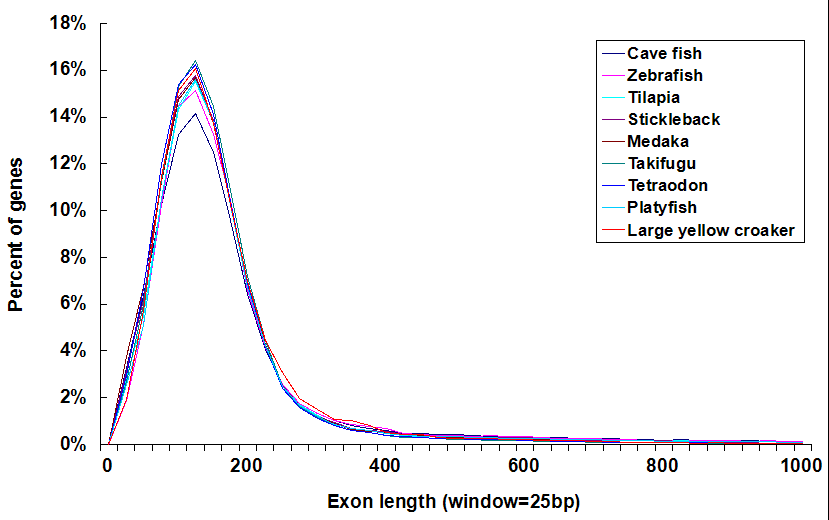

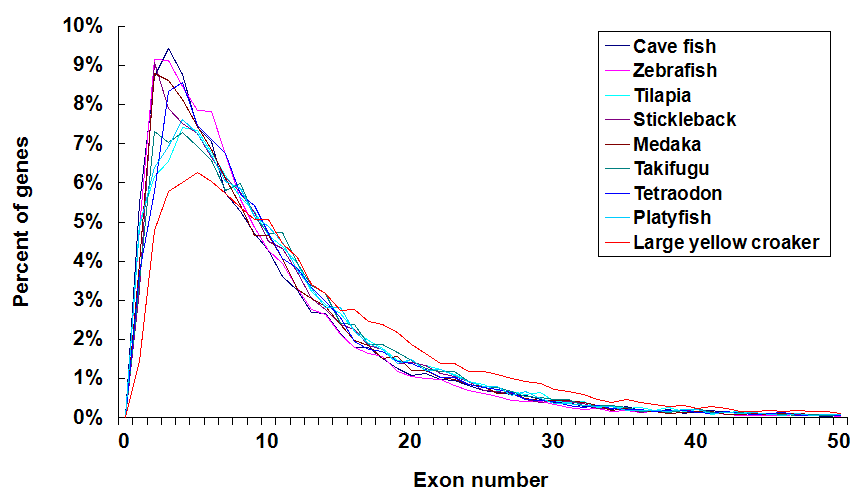

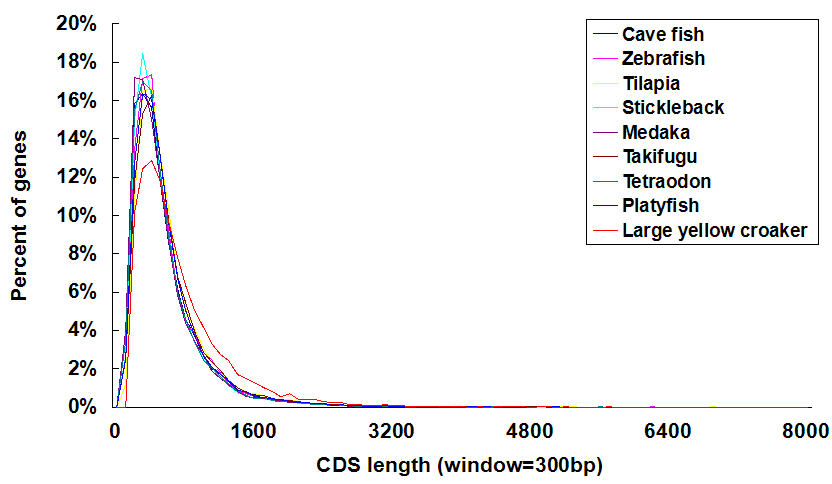

#### FIG. S5. Amplification of 24 novel genes

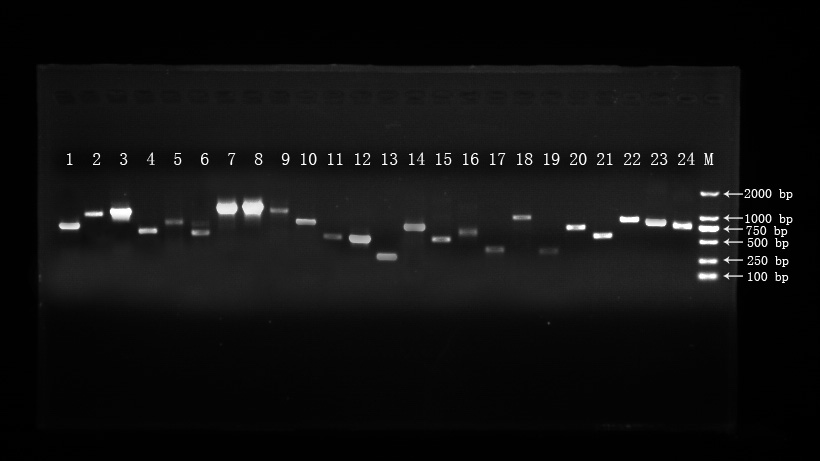

The figure illustrates the amplifications of 24 novel genes predicted in the new assembly but not in the previous two assemblies.

#### FIG. S6. Orthologous profiling among croaker, medaka, stickleback, European sea bass and tetraodon

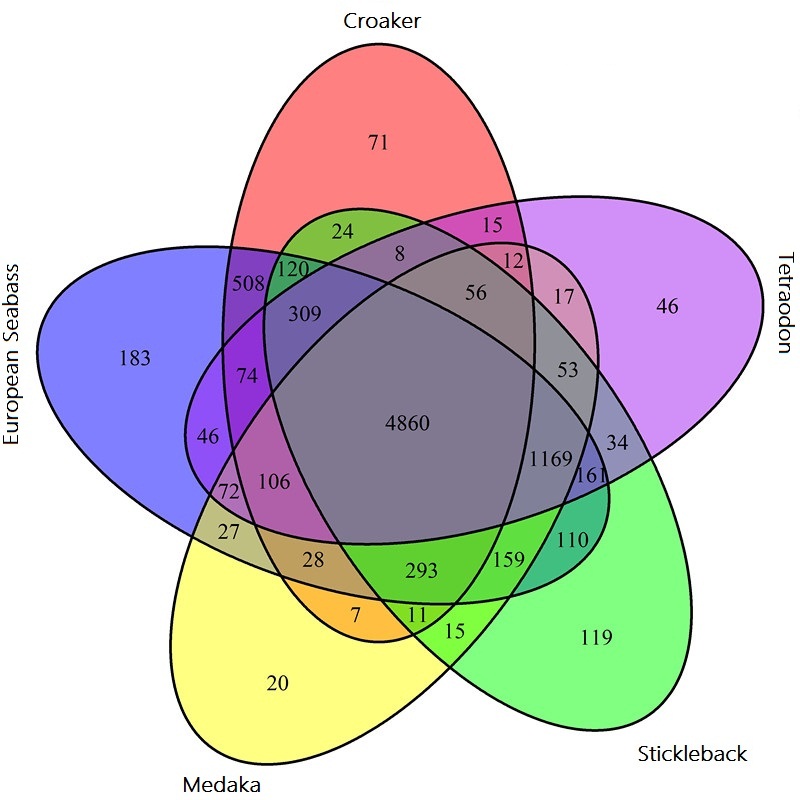

Venn diagram showing unique and overlapping gene families in five close species, including croaker, European Sea bass, medaka, stickleback and tetraodon.

#### FIG. S7. Correlation between intergenic length and genome size and between the proportions of different types of repeats and genome size

(a)

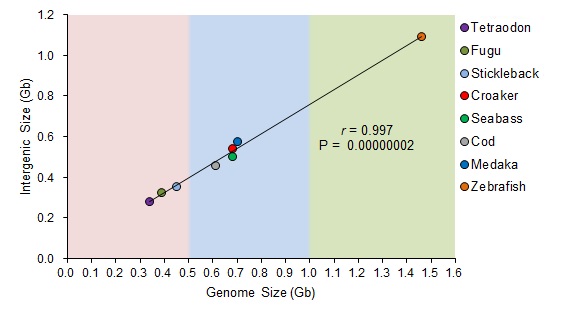

(b)

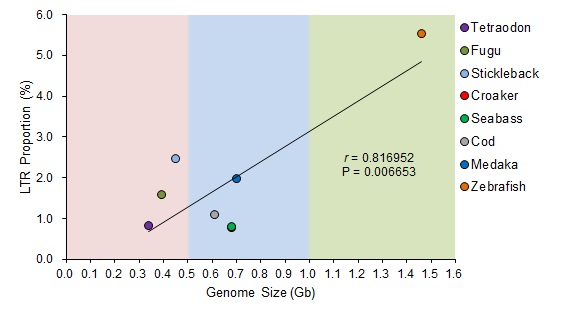

(c)

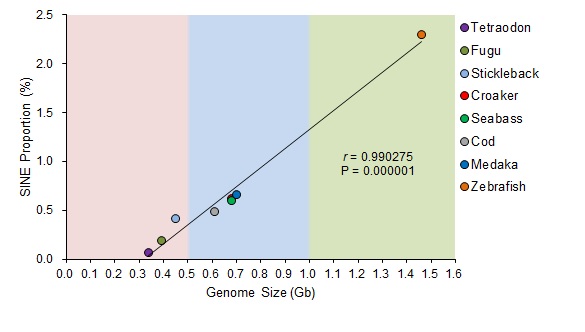

(d)

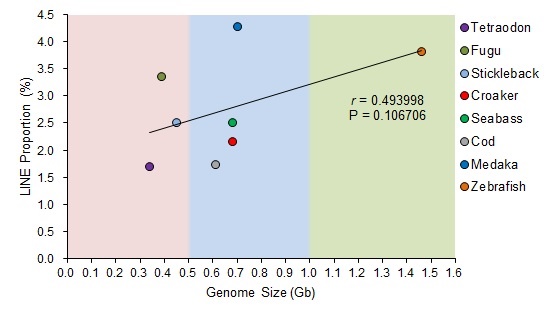

(a) Significant correlation between intergenic length and genome size. (b) Significant correlation between LTR proportion and genome size. (c) Significant correlation between SINE proportion and genome size. (d) Non-significant correlation between LINE proportion and genome size.

#### FIG. S8. Sanger sequencing of two SNPs segregated perfectly with females and males.

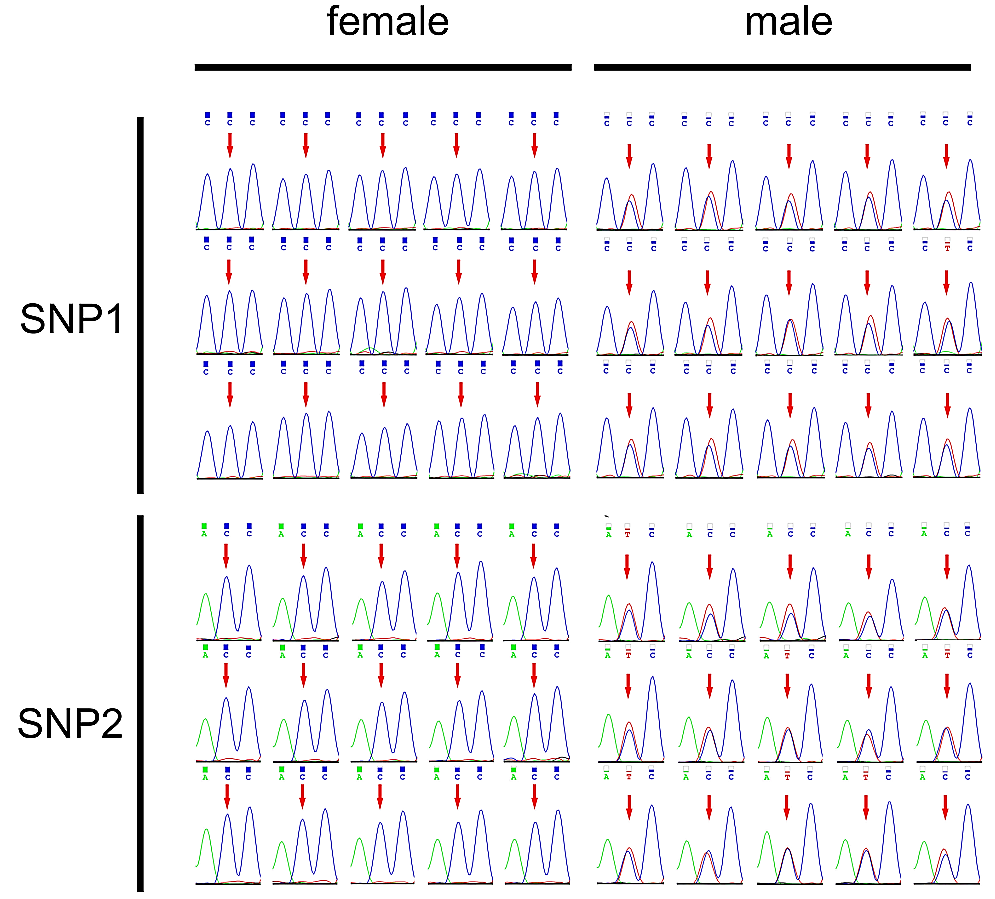

The two SNPs were heterozygous in males and homozygous in females.

#### FIG. S9. K-mer distribution of four individuals

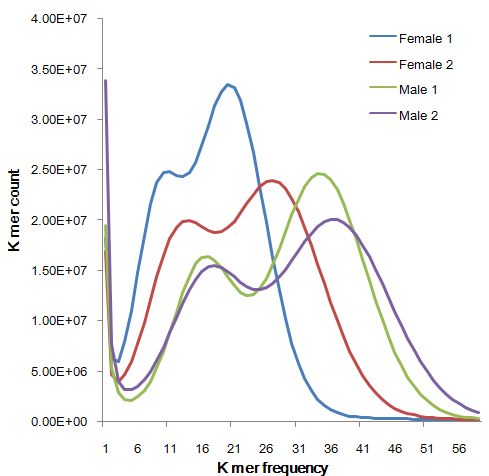

Distribution profile of unique K-mer counts in the cleaned re-sequencing reads of two males and two females. The K-mer size was 17.

#### FIG. S10. The phylogenetic tree of *dmrt* family in vertebrates

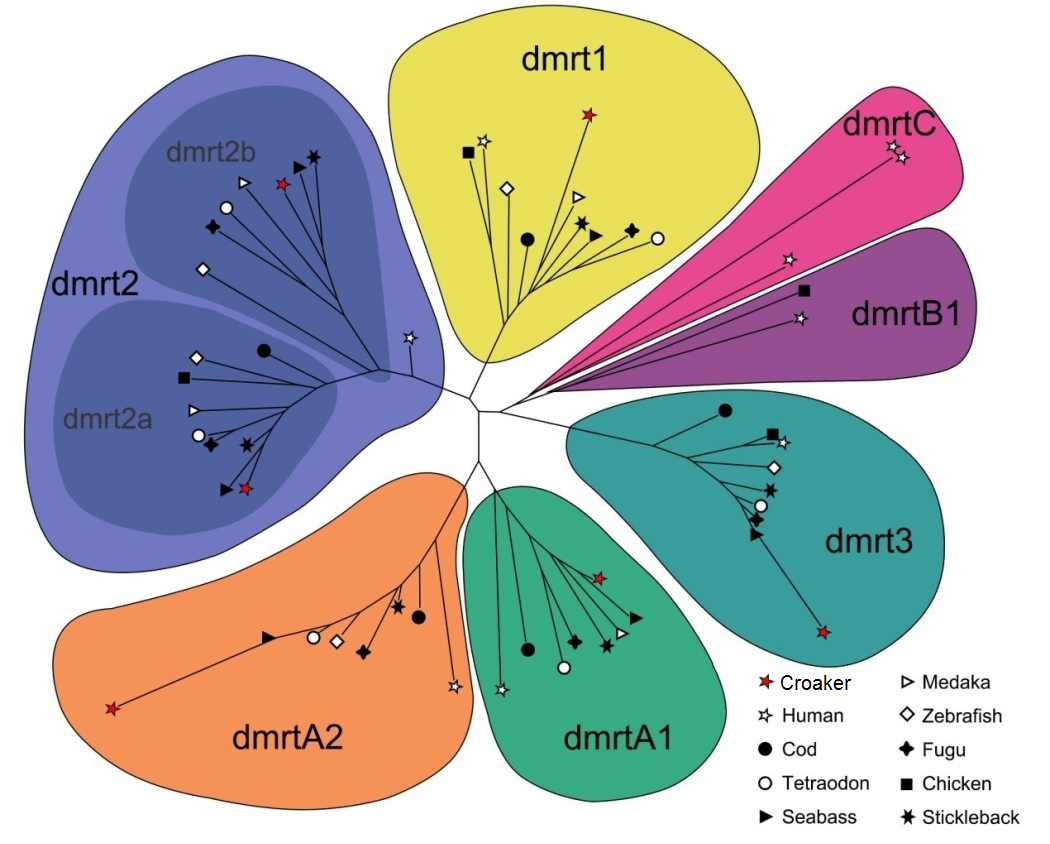

Phylogenetic tree of *dmrt* family in teleosts shows no gene expansion in the croaker genome. The *dmrt* proteins of croaker were marked with a red star.

#### FIG. S11. Profiling of *dmrt1* expression across tissues

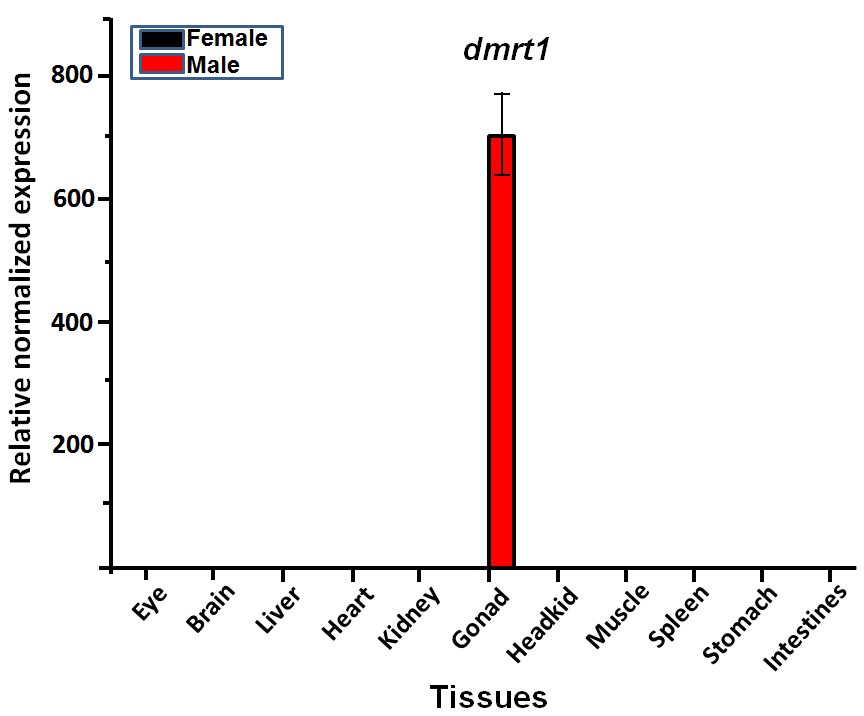

X-axis shows tissues studied; Y-axis shows relative normalized expression level of *dmrt1*. We observed that *dmrt1* was dominantly expressed in male testis.

#### FIG. S12. qRT-PCR validation of two GSD DEGs in males, females and pseudo-males

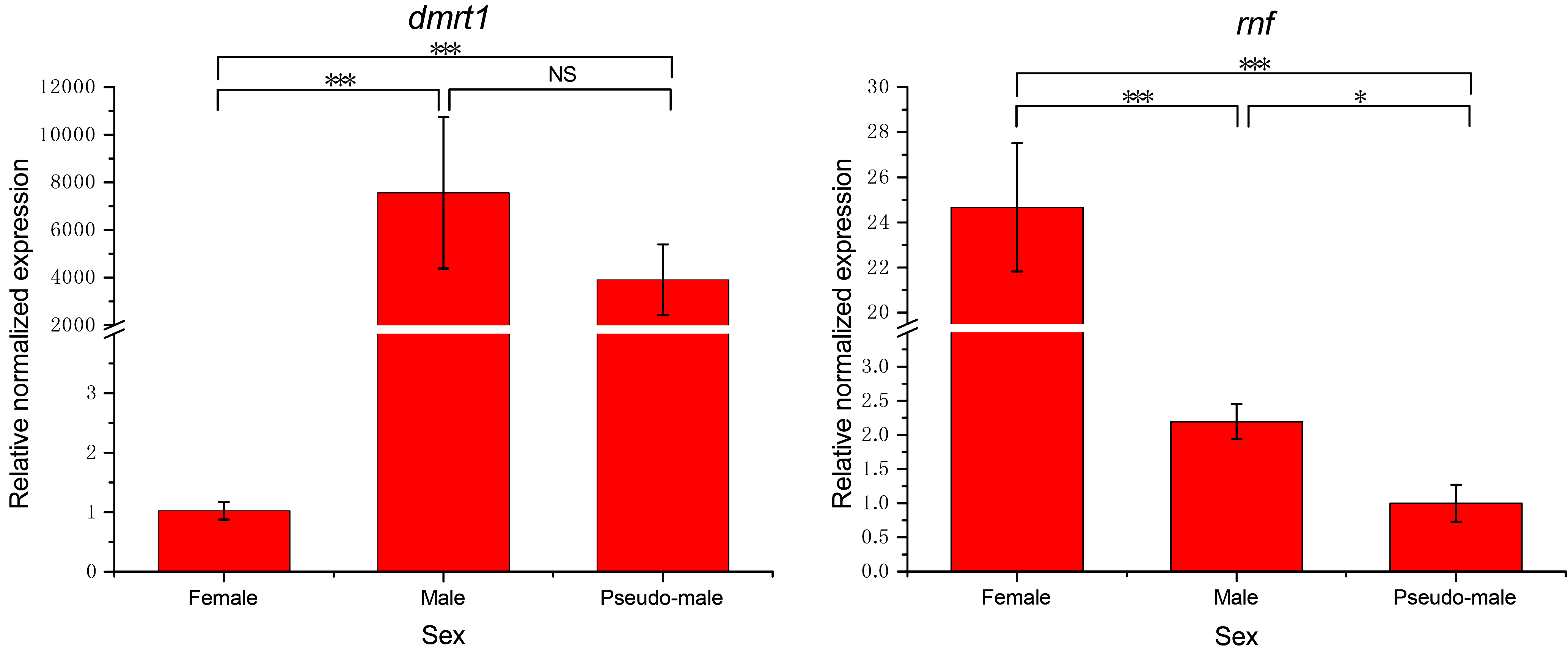

The *dmrt1* and *rnf* in the GSD were predicted to be differentially expressed in GSD and hormone-induced ESR. The *dmrt1* exhibited higher expression level in males and pseudo-males than in females. The *rnf* had higher expression in females than in males and pseudo-males. * means significantly different (*p* < 0.05); *** means extremely significantly different (*p* < 0.001).

#### FIG. S13. Profiling *dmrt1* expression across developmental stages

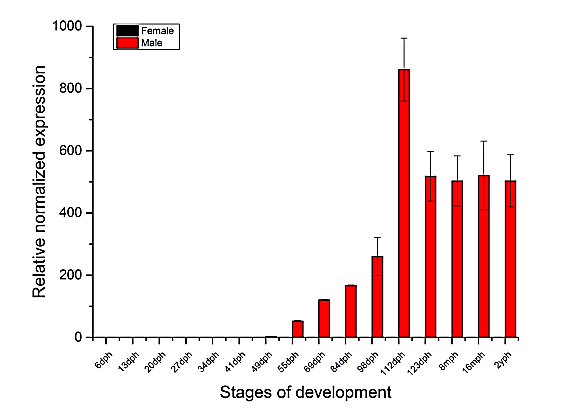

X-axis shows developmental stages interrogated; Y-axis shows relative normalized expression level of *dmrt1*. We observed that the expression of *dmrt1* step-wisely increased along with the maturation of testis.

#### FIG. S14. Sex-biased markers in the transcription factor binding sites of *dmrt1*.

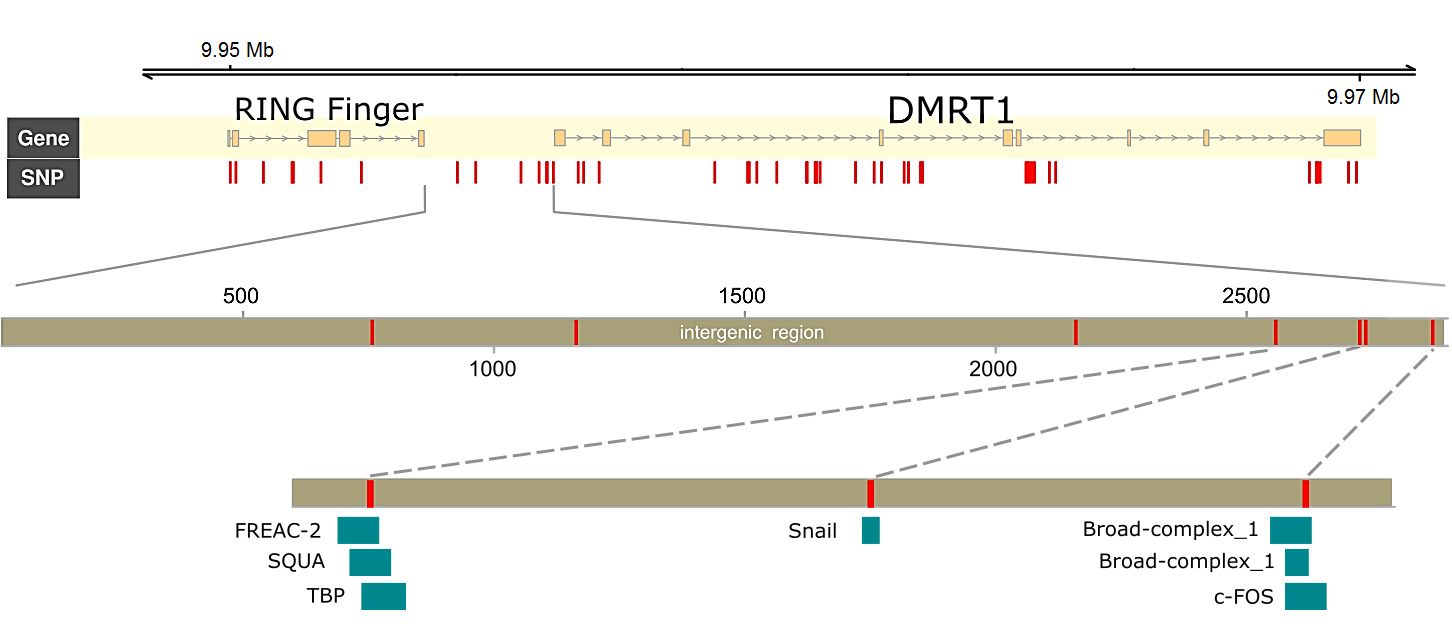

The exonic regions were illuminated by yellow blocks, and sex-related markers were shown by red bars. The markers resided in the 3’UTR of *dmrt1* were depicted in the zoom insets below the gene models, where blue blocks demonstrated the regions for specific transcript factors.

#### FIG. S15. Overlapping of three groups of DEGs.

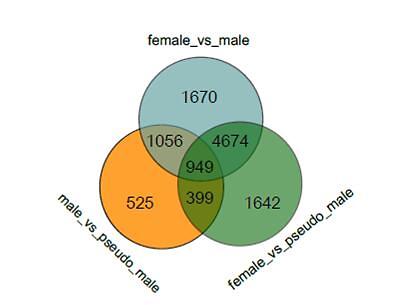

High proportion of DEGs overlapped among female *vs* male, female *vs* pseudo-male, and male *vs* pseudo-male comparisons. Using the DEGs in the comparison of males *vs* pseudo-males as negative control of DEGs, we identified 6,316 hormone-induced DEGs and 6,344 genetic-determining DEGs.

#### FIG. S16. Significantly co-expressed genes with *dmrt1* identified by WGCNA

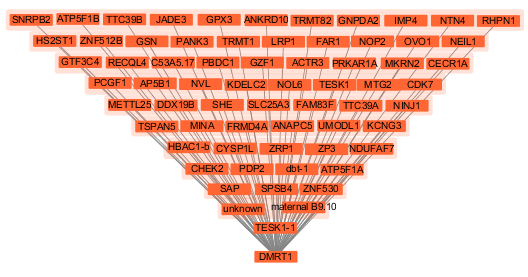

Gene coexpression network including *dmrt1*. 66 genes were identified to have close expression relationship with dmrt1 among three types of gonads using WGCNA with a weight threshold of 0.7. The gene names were shown in the boxes. The lines indicate a close relationship between the co-expressed gens and *dmrt1*. The functions of the 66 genes were described in supplementary table S26.

### Supplementary Tables

#### Table S1. Summary of library construction and sequencing

| **Libraries (insert size)** | **Total Data (Gb)** | **Read length (bp)** | **Sequence Coverage (X)** |
| --- | --- | --- | --- |
| Paired-end (200 bp) | 40.5 | 100 | 57.9 |
| Paired-end (600 bp) | 23.1 | 100 | 33.1 |
| Paired-end (1 kb) | 34.9 | 100 | 49.9 |
| Mate-paired (6 kb) | 16.7 | 100 | 23.9 |
| Mate-paired (8 kb) | 11.2 | 100 | 16.0 |
| Pacbio long reads | 11.9 | 13,661 | 17.5 |
| Total | 126.4 | - | 198.3 |

#### Table S2. Summary of croaker genome assembly

| **Assembly** | **N50 (size/number)** | **N90 (size/number)** | **Longest (kb)** | **Total length** |
| --- | --- | --- | --- | --- |
| Contig | 406.4 kb / 350 | 78.5 kb / 1,919 | 6,173 | 688 Mb |
| Scaffold | 1.23 Mb / 116 | 271 kb / 626 | 9,631 | 689 Mb |
| Chromosomes | 24 chromosomes | | | 510.2 Mb (74%) |

#### Table S3. The quality metrics of previous two assemblies and our assembly

|  | **The assembly**  **of Wu *et al.*** | **The assembly**  **of Ao *et al.*** | **The assembly**  **in our study** |
| --- | --- | --- | --- |
| Contig N50 | 25.7 kb | 63.11 kb | 406.4 kb |
| Scaffold N50 | 498.7 kb | 1.03 Mb | 1.23 Mb |
| Genome length without gaps | 618.9 Mb | 661.3 Mb | 688 Mb |
| Gene number | 19,362 | 25,401 | 23,272 |
| Exon number | 186,960 | 251,617 | 313,286 |

#### Table S4. Mapping ratio of Illumina genomic sequencing reads to our croaker assembly

| **Libraries** | **Clean pairs** | **Mapped pairs** | **Mapping ratio (%)** |
| --- | --- | --- | --- |
| Illumina 200 bp | 375,657,569 | 342,306,096 | 91.12 |
| Illumina 600 bp | 221,534,792 | 199,958,845 | 90.26 |
| Illumina 1 kb | 334,528,394 | 304,531,885 | 91.03 |
| Illumina 6 kb | 166,057,313 | 136,953,373 | 82.47 |
| Illumina 8 kb | 105,853,185 | 94,185,743 | 88.98 |
| Total | 1,203,631,253 | 1,077,935,942 | 89.56 |

#### Table S5. Mapping ratio of re-sequencing reads to our croaker assembly

|  | **Female population** | **Male population** | **Female 1** | **Female 2** | **Male 1** | **Male 2** |
| --- | --- | --- | --- | --- | --- | --- |
| Read number | 203,335,896 | 456,819,930 | 202,840,387 | 256,910,035 | 321,325,895 | 341,405,482 |
| Data (Gb) | 20.34 | 45.68 | 20.28 | 25.69 | 32.13 | 34.14 |
| Mapping ratio (%) | 92.96 | 92.39 | 91.94 | 92.7 | 92.42 | 92.42 |

#### Table S6. Mapping ratio of CEGMA proteins to our croaker assembly

|  | **CEG proteins** | **Mapped proteins** | **Completeness** |
| --- | --- | --- | --- |
| Total | 248 | 246 | 99.2% |
| Group 1 | 66 | 65 | 98.5% |
| Group 2 | 56 | 55 | 98.2% |
| Group 3 | 61 | 61 | 100% |
| Group 4 | 65 | 65 | 100% |

#### Table S7. Summary of croaker genome components

|  | **Prediction** | **Number** | **Total Length**  **(Mb)** | **Percentage of**  **Genome (%)** |
| --- | --- | --- | --- | --- |
| Repeat | *ab initio* repeat prediction | 978,286 | 98.3 | 14.27 |
|  | homolog based repeat prediction | 6,731 | 0.66 | 0.10 |
| Gene | gene | 23,272 | 312.4 | 45.3 |
|  | exon | 313,286 | 49.57 | 7.2 |

#### Table S8. The statistics of gene structures of croaker and other teleosts

| **Species** | **Genome**  **size (Mb)** | **No. genes** | **Mean CDS**  **length (bp)** | **Median**  **CDS (bp)** | **Mean intergenic region (bp)** | **Median intergenic region（bp）** | **No. exons**  **per gene** | **Mean exon size(bp)** | **Mean intron size(bp)** | **Median intron size（bp）** |
| --- | --- | --- | --- | --- | --- | --- | --- | --- | --- | --- |
| *L. crocea* | 689 | 23,125 | 2,066 | 1,572 | 44,405 | 20,029 | 13.41 | 158 | 872 | 341 |
| *D. rerio* | 1,464 | 25,465 | 1634 | 1,209 | 81,334 | 23,931 | 9.92 | 240 | 3,220 | 1,225 |
| *G. morhua* | 608 | 20,095 | 1458 | 1,074 | 28,941 | 11,620 | 12.75 | 114 | 1,283 | 414 |
| *G. aculeatus* | 447 | 20,787 | 1547 | 1,146 | 33,580 | 12,038 | 10.69 | 159 | 907 | 413 |
| *O. latipes* | 700 | 19,699 | 1514 | 1,122 | 68,063 | 22,658 | 10.46 | 154 | 1,332 | 457 |
| *D. labrax* | 680 | 26,719 | 1566 | 1,146 | 38,889 | 14,474 | 9.10 | 292 | 1,321 | 365 |
| *T. rubripes* | 393 | 18,523 | 1692 | 1,287 | 26,531 | 10,408 | 11.52 | 145 | 700 | 348 |
| *T. nigroviridis* | 342 | 19,602 | 1515 | 1,143 | 29,613 | 10,936 | 10.64 | 148 | 620 | 320 |
| *G. gallus* | 1,073 | 15,508 | 1668 | 1,230 | 107,786 | 27,821 | 10.91 | 234 | 2,831 | 962 |
| *H. sapiens* | 3,547 | 19,826 | 1722 | 1,281 | 254,516 | 59,302 | 10.84 | 283 | 6,007 | 1,726 |

#### Table S9. Redundant genes in the previous assemblies

|  | **Query Accession** | **Target**  **Accession** | **Identity** | **Alignment length** | **Mismatches** | **Gap openings** | **Query**  **Start** | **Query**  **End** | **Target**  **Start** | **Target**  **End** | **E-value** | **Score** | **Query**  **Length (aa)** | **Target**  **Length**  **(aa)** |
| --- | --- | --- | --- | --- | --- | --- | --- | --- | --- | --- | --- | --- | --- | --- |
| **the assembly**  **of Ao *et al.*** | KKF08654.1 | KKF28188.1 | 100.00 | 319 | 0 | 0 | 1 | 319 | 1 | 319 | 0.0 | 669 | 319 | 319 |
|  | KKF08688.1 | KKF28512.1 | 100.00 | 146 | 0 | 0 | 518 | 663 | 1 | 146 | 1e-102 | 313 | 663 | 146 |
|  | KKF08734.1 | KKF29731.1 | 100.00 | 435 | 0 | 0 | 50 | 484 | 1 | 435 | 0.0 | 907 | 484 | 435 |
|  | KKF08754.1 | KKF17037.1 | 100.00 | 357 | 0 | 0 | 1 | 357 | 1 | 357 | 0.0 | 725 | 357 | 357 |
|  | KKF08799.1 | KKF08801.1 | 100.00 | 49 | 0 | 0 | 1 | 49 | 145 | 193 | 3e-27 | 98.6 | 49 | 193 |
|  | KKF08821.1 | KKF08820.1 | 100.00 | 152 | 0 | 0 | 1 | 152 | 966 | 1117 | 4e-106 | 330 | 152 | 1117 |
|  | KKF08948.1 | KKF30021.1 | 99.18 | 610 | 5 | 0 | 1 | 610 | 6 | 615 | 0.0 | 1231 | 610 | 615 |
|  | KKF08833.1 | KKF23486.1 | 100.00 | 262 | 0 | 0 | 1 | 262 | 1 | 262 | 0.0 | 551 | 262 | 262 |
| **the assembly**  **of Wu *et al.*** | NP_001290263 | XP_019112549 | 100.00 | 186 | 0 | 0 | 1 | 186 | 1 | 186 | 5e-120 | 337 | 186 | 186 |
|  | NP_001306868 | XP_019130461 | 100.00 | 318 | 0 | 0 | 1 | 318 | 1 | 318 | 0.0 | 662 | 318 | 318 |
|  | XP_010727503 | XP_019122672 | 100.00 | 179 | 0 | 0 | 1 | 179 | 1 | 179 | 7e-134 | 371 | 179 | 179 |
|  | XP_010727505 | XP_019128772 | 100.00 | 369 | 0 | 0 | 1 | 369 | 1 | 369 | 0.0 | 716 | 369 | 369 |
|  | XP_010727506 | XP_019122653 | 100.00 | 373 | 0 | 0 | 1 | 373 | 1 | 373 | 0.0 | 744 | 373 | 373 |
|  | XP_010727549 | XP_019122684 | 100.00 | 211 | 0 | 0 | 1 | 211 | 1 | 211 | 1e-138 | 386 | 211 | 211 |
|  | XP_010727564 | XP_019134055 | 100.00 | 455 | 0 | 0 | 1 | 455 | 1 | 455 | 0.0 | 824 | 455 | 455 |
|  | XP_010727601 | XP_019109162 | 100.00 | 409 | 0 | 0 | 1 | 409 | 1 | 409 | 0.0 | 827 | 409 | 409 |

The predicted genes of previous two assemblies were downloaded from NCBI Assembly database. The longest protein of each gene was selected to represent the gene. Then we performed an all-against-all Blastp search on the predicted genes from each of previous two assemblies.

#### Table S10. Annotations of croaker genes against known protein databases

| **Database** | **Number** | **Percent (%)** |
| --- | --- | --- |
| Swiss-Prot | 13,706 | 58.90 |
| TrEMBL | 21,285 | 91.46 |
| NR database | 21,303 | 91.54 |
| GO | 18,470 | 79.36 |
| KEGG pathway | 12,010 | 51.61 |
| Total | 23,033 | 98.97 |

#### Table S11. Primers for amplification of twenty-four randomly selected novel genes

| **Gene Symbol** | **Sequence name** | **Primer sequence (5′-3′)** | **Annealing temperature** |
| --- | --- | --- | --- |
| *rars2* | JMU_LC_G_04383 | F: AAGTTGAAGCAGGACAGTGTG | 60℃ |
|  |  | R: GGCGAGATCTCTGGTGATGT |  |
| *arg2-b* | JMU_LC_G_10671 | F: CAGCAACACCTCTCAGAGCTT | 60℃ |
|  |  | R: GATCGGTCGCTGCTTTCT |  |
| *gpr34* | JMU_LC_G_04298 | F: TCAACAACCCTCAGACGCGAT | 60℃ |
|  |  | R: AGCTGGGACAGGATGTAGAAT |  |
| *chmp1b* | JMU_LC_G_13416 | F: CCGTGCGTGAGGACTTGT | 60℃ |
|  |  | R: TCCCACTGATCCTGTCTGAC |  |
| *slc5a9* | JMU_LC_G_04590 | F: GTTCCCAGATGAAGTAGGTT | 60℃ |
|  |  | R:GGGCTGAGGTAGAGTTGACT |  |
| *eif2b1* | JMU_LC_G_10610 | F:GATGCAGCTGTGGGGTACGT | 61℃ |
|  |  | R:TAAAGCTTGATGAGTTCGTCGCT |  |
| *mgat2* | JMU_LC_G_10340 | F:AGAAGCGACAAATGACACGGT | 58℃ |
|  |  | R:AATCTGGGCAGCTGCTCTT |  |
| *arg2* | JMU_LC_G_01219 | F:TTCCGTCTTCTCAGGGCT | 58℃ |
|  |  | R:CAACCCAAATGAGACACAGGT |  |
| *uspl1* | JMU_LC_G_13054 | F:GCTTGCAGTGACATGGTAA | 57℃ |
|  |  | R: AGTCTCGTCCCCATTGGT |  |
| *c7orf43* | JMU_LC_G_16038 | F:TTGTGCCAATCTCCCCTC | 55℃ |
|  |  | R:TAGCTGCCATCACTCCAT |  |
| *dkk2* | JMU_LC_G_16248 | F:GCTGTTGGTGCTTTGTGT | 54℃ |
|  |  | R: AGATGCGAGTCCAGAAGT |  |
| *ndufb4* | JMU_LC_G_16902 | F:CGCTGGACCCTAATGAGTAT | 54℃ |
|  |  | R:AATCTGGGCCTCCTTCCTAT |  |
| *gvin1* | JMU _LC_G_19422 | F:AAGGATACGCAGTCAGTGAAGAT | 58℃ |
|  |  | R:AGTTGTAGAGATTGCCTGGTAC |  |
| *ack1* | JMU _LC_G_23103 | F:TCCCTCAGAAGTTCATGC | 53℃ |
|  |  | R:ACAAACCCACTGCCTCAT |  |
| *si:ch73-30l9.1* | JMU _LC_G_22615 | F:GATGTGCACTCTGAGTCTTT | 53℃ |
|  |  | R: AAGAGAGATGCTGCTTGTCT |  |
| *grin2* | JMU _LC_G_22605 | F:CCACACAGGCAATATACCT | 55℃ |
|  |  | R:TTAGGAGGAGGATGGCTAT |  |
| *nhe3b* | JMU _LC_G_21835 | F:TGAACTTGAAGGTTGACG | 53℃ |
|  |  | R: GATGATTGATAGAGCGCAT |  |
| *zcchc3* | JMU _LC_G_21820 | F:GTGACTACGTAGGTGAACATCT | 55℃ |
|  |  | R: CACATAAGCTGCAGACACT |  |
| *ache* | JMU _LC_G_21782 | F:ACACAGAGGGAGGTATGGT | 57℃ |
|  |  | R: GAACTGTTATGTCCTCCTCAG |  |
| *kiaa1586* | JMU _LC_G_21623 | F:CAGCGGGTCATTCAAACT | 56℃ |
|  |  | R: GGCCTCGAAGACTCTGATCT |  |
| *sat1* | JMU _LC_G_21494 | F:ATGTCTGACCAGGTGAAGAT | 53℃ |
|  |  | R: TCAAAGCGTATGAAGTGC |  |
| *NA* | JMU _LC_G_21156 | F:CGCATCCAGGCTTCAGTT | 55℃ |
|  |  | R: CGTGTAATGCGGGTGAAAT |  |
| *diablo* | JMU _LC_G_18935 | F:GGAGAGGAGCAGCATGTAT | 55℃ |
|  |  | R: TCAGTTTCCGTGCCTCCT |  |
| *NA* | JMU _LC_G_16992 | F:GCACACACACACCATGAGT | 59℃ |
|  |  | R:GGACTCTCTGACCTGGATG |  |

NA indicates no homologs in other species.

#### Table S12. Assignment ratio of teleost genes to TreeFam families

| **Species** | **Assigned genes** | **TreeFam Families** | **Assignment Ratio** |
| --- | --- | --- | --- |
| *L. crocea* | 18,979 | 6,960 | 0.815 |
| *D. labrax* | 22,647 | 8225 | 0.847 |
| *D. rerio* | 16,962 | 7,640 | 0.666 |
| *O. latipes* | 12,706 | 6,905 | 0.645 |
| *G. aculeatus* | 14,651 | 7,501 | 0.705 |
| *T. rubripes* | 13,377 | 7,022 | 0.722 |
| *T. nigroviridis* | 13,566 | 7,038 | 0.692 |
| *G. morhua* | 13,150 | 7,005 | 0.654 |

**Table S13. Systematic cross-species comparative analysis**

|  | **1：1：1** | **x：x：x** | **Fish specific** | **Others** | **Total** |
| --- | --- | --- | --- | --- | --- |
| *D.rerio* | 877 | 9,742 | 288 | 6,055 | 16,962 |
| *L. crocea* | 877 | 9,349 | 149 | 6,476 | 16,851 |
| *G.morhua* | 877 | 8,425 | 124 | 3,724 | 13,150 |
| *G. aculeatus* | 877 | 8,939 | 184 | 4,651 | 14,651 |
| *O.latipes* | 877 | 8,208 | 166 | 3,455 | 12,706 |
| *D. labrax* | 877 | 10,323 | 344 | 11,103 | 22,647 |
| *T. rubripes* | 877 | 8,754 | 148 | 3,598 | 13,377 |
| *T. nigroviridis* | 877 | 8,795 | 137 | 3,757 | 13,566 |
| *G. gallus* | 877 | 6,310 | - | 3,887 | 11,074 |
| *H.sapiens* | 877 | 7,519 | - | 6,700 | 15,096 |

'1:1:1' indicates universal single-copy genes. 'X:X:X' indicates orthologs exist in all genomes (missing in one species not allowed), with 'X' meaning one or more orthologs per species. Those ‘1:1:1’ genes were not included in ‘X:X:X’. 'Fish specific' indicates orthologs exist in teleost genomes but not in human and chicken genome. ‘Others’ indicates genes which are assigned one gene family but do not fit into the categories of ‘1:1:1’, ‘X:X:X’ and ‘Fish specific’ includes genes without homologs to other species.

#### Table S14. Repeat content in croaker genome

| **Repeat Elements** | **Copies** | | **Bases** | | **Percent (%)** |
| --- | --- | --- | --- | --- | --- |
| **Interspersed repeats** |  |  |  |  |  |
| SINE | 30,934 |  | 4,216,243 |  | 0.61 |
| LINE | 72,100 |  | 14,636,934 |  | 2.12 |
| LTR | 18,860 |  | 5,370,566 |  | 0.78 |
| DNA | 183,679 |  | 30,133,485 |  | 4.37 |
| Unclassified | 324,753 |  | 45,782,755 |  | 6.64 |
| Subtotal | 630,326 |  | 100,139,983 |  | 14.53 |
| **Tandem repeats** |  |  |  |  |  |
| Satellites | 840 |  | 151,626 |  | 0.022 |
| Simple repeats | 385,568 |  | 18,201,047 |  | 2.64 |
| Low complexity | 42,488 |  | 2,447,859 |  | 0.36 |
| Subtotal | 428,896 |  | 20,800,532 |  | 3.02 |
| **Small RNA** | 10,472 |  | 1,280,462 |  | 0.19 |
| **Total** | 1,069,694 |  | 122,220,977 |  | 17.74 |

#### Table S15. Interspersed repeat components of eight teleost genomes

|  | **SINE bases** | | **LINE bases** | | **LTR bases** | | **DNA transposon bases** | | **Unclassified repeat bases** | | **Total bases** |
| --- | --- | --- | --- | --- | --- | --- | --- | --- | --- | --- | --- |
|  | De novo | Homolog  based | De novo | Homolog  based | De novo | Homolog  based | De novo | Homolog  based | De novo | Homolog  based |  |
| *D.rerio* | 31,400,055 | 126,866 | 51,347,876 | 994,607 | 75,170,996 | 671,700 | 549,672,546 | 6,627,643 | 24,333,808 | 233,960 | 740,580,057 |
| *L. crocea* | 4,168,323 | 47,920 | 12,130,820 | 2,506,114 | 3,004,297 | 2,366,269 | 27,755,813 | 2,377,672 | 45,602,066 | 180,689 | 100,139,983 |
| *G.morhua* | 4,002,218 | 37,366 | 12,245,209 | 2,089,256 | 5,615,580 | 3,413,707 | 41,316,723 | 2,706,718 | 50,552,342 | 116,952 | 122,096,071 |
| *G. aculeatus* | 1,884,234 | 13,271 | 10,944,429 | 635,371 | 8,994,124 | 2,428,446 | 19,453,103 | 1,540,083 | 16,995,110 | 82,386 | 62,970,557 |
| *O.latipes* | 5,724,779 | 26,089 | 34,043,464 | 3,210,345 | 14,350,447 | 2,908,987 | 88,927,243 | 2,832,270 | 90,807,466 | 126,768 | 242,957,858 |
| *D. labrax* | 4,053,469 | 57,032 | 14,358,746 | 2,655,223 | 2,504,563 | 2,901,089 | 3,284,7124 | 2,969,782 | 66,192,385 | 218,545 | 128,757,958 |
| *T. rubripes* | 737,794 | 28,871 | 12,141,972 | 986,873 | 4,942,640 | 1,222,419 | 6,142,506 | 1,371,823 | 5,507,286 | 103,026 | 3,3185,210 |
| *T. nigroviridis* | 232,688 | 22,682 | 5,685,306 | 432,086 | 2,636,770 | 338,900 | 4,444,735 | 641,868 | 9,178,825 | 100,575 | 23,714,435 |

We identified interspersed repeat using a two-step approach. Each genome was firstly masked using these *de novo* repeat libraries and then further searched for homologous repeats against RepBase fish library.

#### Table S16. Six sex-linked markers in the integrated chr9 of croaker

| **Linkage group** | **SNP marker*** | **Genetic**  **position (cM)** | **LOD** | **GWAS**  **P value** | **Heteromorphic male numbers** | **Homomorphic**  **Male number** | **Heteromorphic**  **female numbers** | **Homomorphic**  **female number** |
| --- | --- | --- | --- | --- | --- | --- | --- | --- |
| 9 | C38623853_152 | 29.629 | 45.68 | 2.74e-08 | 36 | 1 | 0 | 35 |
| 9 | scaffold345633_271 | 31.622 | 37.7 | 2.42e-06 | 31 | 6 | 2 | 33 |
| 9 | C38780343_116 | 31.818 | 38.42 | 1.80e-07 | 35 | 2 | 1 | 34 |
| 9 | C39250574_131 | 32.842 | 52.68 | 2.26e-07 | 36 | 1 | 2 | 33 |
| 9 | scaffold365802_91 | 35.034 | 46.74 | 4.56e-07 | 36 | 1 | 3 | 32 |
| 9 | C39162591_196 | 35.076 | 47.84 | 9.55e-07 | 0 | 37 | 26 | 9 |

*The genetic positions of these markers were available at our published genetic map.

#### Table S17. 617 sites with polymorphism segregation in two sexes

(in a separate excel file)

#### Table S18. The functions of 79 genes in croaker GSD locus

(in a separate excel file)

#### Table S19. Primers used to validate two sex-biased markers.

| **Genome region** | **Marker** | **F Primer** | **R Primer** |
| --- | --- | --- | --- |
| chr9: 9951961 | SNP1 | ATCTGTCAACCACTGTATCATCTG | GGATGGCGTTTGGCTGAG |
| chr9: 9950079 | SNP2 | GCATTTCCTCGTTCGTTATTCAG | CCGCTCTTACCGTTCACAATC |

#### Table S20. Estimated genome sizes of four individuals

| **Sample** | **Sequencing bases (Gb)** | **Estimated genome size (Mb)** |
| --- | --- | --- |
| Female 1 | 20.28 | 730.8 |
| Female 2 | 25.69 | 726.7 |
| Male 1 | 32.13 | 727.4 |
| Male 2 | 34.14 | 730.5 |

We estimated the genome sizes of four samples with each short-insert paired-end library.

#### Table S21. The sex-biased SNPs and Ks of genes in the GSD locus

| **chr** | **SNP locus** | **gene** | **Gene Symbol** | **exon begin** | **exon end** | **mutation_impact_type** | **mutation_function** | **Ks** |
| --- | --- | --- | --- | --- | --- | --- | --- | --- |
| chr9 | 7191575 | JMU_LC_G_00780 | *csn1* | 7191507 | 7191576 | synonymous | gaC/gaT\|D426 | 0.0020 |
| chr9 | 7284125 | JMU_LC_G_00757 | *LOC104930121* | 7284029 | 7284636 | nonsynonymous | cCg/cTg\|P245L | 0.0000 |
| chr9 | 7528732 | JMU_LC_G_00776 | *parp4* | 7528527 | 7528842 | stop loss | tAg/tGg\|*69W | 0.0000 |
| chr9 | 8013982 | JMU_LC_G_00784 | *ift88* | 8013935 | 8013988 | synonymous | gcT/gcA\|A117 | 0.0016 |
| chr9 | 9029791 | JMU_LC_G_02402 | *cdca2* | 9028750 | 9030004 | nonsynonymous | Tca/Cca\|S426P | 0.0000 |
| chr9 | 9050159 | JMU_LC_G_02407 | *sema4c* | 9050142 | 9050232 | synonymous | gtC/gtT\|V559 | 0.0001 |
| chr9 | 9050165 | JMU_LC_G_02407 | *sema4c* | 9050142 | 9050232 | synonymous | aaA/aaG\|K557 |  |
| chr9 | 9566799 | JMU_LC_G_02411 | *txnrd2* | 9566655 | 9566833 | synonymous | gaA/gaG\|E241 | 0.0056 |
| chr9 | 8483521 | JMU_LC_G_02418 | *chfr* | 8483456 | 8483608 | stop gain | Gga/Tga\|G324* | 0.0042 |
| chr9 | 8483756 | JMU_LC_G_02418 | *chfr* | 8483743 | 8483902 | nonsynonymous | aTg/aCg\|M290T |  |
| chr9 | 8501322 | JMU_LC_G_02432 | *fbrsl1* | 8501212 | 8502457 | nonsynonymous | Cgt/Tgt\|R1149C | 0.0000 |
| chr9 | 9950081 | JMU_LC_G_02421 | *LOC104934771* | 9949992 | 9950129 | nonsynonymous | Cct/Tct\|P45S | 0.0000 |
| chr9 | 9951962 | JMU_LC_G_02421 | *LOC104934771* | 9951665 | 9952283 | nonsynonymous | cCc/cTc\|P160L |  |

#### *Table S22. The predicted biological functions of the sex-biased SNPs and InDels around dmrt1*

| **Chr** | **Location** | **Base in females** | **Base in males** | **Function** | **Related gene** | **Gene Symbols** |
| --- | --- | --- | --- | --- | --- | --- |
| chr9 | 9955386 | C | A | UTR_3_PRIME | JMU_LC_G_02421 | *rnf* |
| chr9 | 9956386 | C | G | UTR_3_PRIME | JMU_LC_G_02421 | *rnf* |
| chr9 | 9956954 | A | G | UTR_3_PRIME | JMU_LC_G_02421 | *rnf* |
| chr9 | 9956966 | TCC | T | UTR_3_PRIME | JMU_LC_G_02421 | *rnf* |
| chr9 | 9957100 | C | T | UTR_5_PRIME | JMU_LC_G_02420 | *dmrt1* |
| chr9 | 9957651 | G | T | INTRON | JMU_LC_G_02420 | *dmrt1* |
| chr9 | 9957773 | G | A | INTRON | JMU_LC_G_02420 | *dmrt1* |
| chr9 | 9958114 | A | G | INTRON | JMU_LC_G_02420 | *dmrt1* |
| chr9 | 9960672 | G | T | INTRON | JMU_LC_G_02420 | *dmrt1* |
| chr9 | 9961399 | CA | C | INTRON | JMU_LC_G_02420 | *dmrt1* |
| chr9 | 9961434 | C | T | INTRON | JMU_LC_G_02420 | *dmrt1* |
| chr9 | 9962044 | C | T | INTRON | JMU_LC_G_02420 | *dmrt1* |
| chr9 | 9962702 | G | A | INTRON | JMU_LC_G_02420 | *dmrt1* |
| chr9 | 9962715 | G | T | INTRON | JMU_LC_G_02420 | *dmrt1* |
| chr9 | 9962907 | T | TTG | INTRON | JMU_LC_G_02420 | *dmrt1* |
| chr9 | 9962928 | G | T | INTRON | JMU_LC_G_02420 | *dmrt1* |
| chr9 | 9963001 | T | C | INTRON | JMU_LC_G_02420 | *dmrt1* |
| chr9 | 9964361 | TAAATGAGTTTCACTC | T | INTRON | JMU_LC_G_02420 | *dmrt1* |
| chr9 | 9964954 | C | G | INTRON | JMU_LC_G_02420 | *dmrt1* |
| chr9 | 9965222 | C | T | INTRON | JMU_LC_G_02420 | *dmrt1* |
| chr9 | 9968070 | G | A | INTRON | JMU_LC_G_02420 | *dmrt1* |
| chr9 | 9968212 | C | T | INTRON | JMU_LC_G_02420 | *dmrt1* |
| chr9 | 9973821 | C | G | INTRON | JMU_LC_G_02420 | *dmrt1* |
| chr9 | 9974060 | C | T | INTRON | JMU_LC_G_02420 | *dmrt1* |
| chr9 | 9974862 | A | C | UTR_3_PRIME | JMU_LC_G_02420 | *dmrt1* |
| chr9 | 9975252 | G | A | UTR_3_PRIME | JMU_LC_G_02420 | *dmrt1* |
| chr9 | 9975254 | G | T | UTR_3_PRIME | JMU_LC_G_02420 | *dmrt1* |
| chr9 | 9975255 | AGC | A | UTR_3_PRIME | JMU_LC_G_02420 | *dmrt1* |
| chr9 | 9975262 | A | T | UTR_3_PRIME | JMU_LC_G_02420 | *dmrt1* |
| chr9 | 9975264 | A | G | UTR_3_PRIME | JMU_LC_G_02420 | *dmrt1* |
| chr9 | 9975265 | A | C | INTERGENIC |  |  |
| chr9 | 9975319 | GTGT | G | INTERGENIC |  |  |
| chr9 | 9977316 | G | T | INTERGENIC |  |  |
| chr9 | 9977594 | A | G | INTERGENIC |  |  |

UTR_3_PRIME and UTR_5_PRIME mean the 3’UTR and 5’UTR region, respectively.

#### Table S23. Cleaned RNA-seq reads of male, pseudo-male and female gonads of croaker

| **Sample** | **Reads** | **Average length (bp)** | **Total bases** |
| --- | --- | --- | --- |
| male 3 (M3) | 35,309,772 | 121.8 | 4,300,730,230 |
| male 4 (M4) | 30,987,356 | 122 | 3,780,457,432 |
| male 5 (M5) | 31,519,684 | 121.8 | 3,839,097,511 |
| pesudo-male 1 (PM1) | 34,653,950 | 121.8 | 4,220,851,110 |
| pesudo-male 2 (PM2) | 28,931,590 | 121.9 | 3,526,760,821 |
| pesudo-male 3 (PM3) | 36,766,490 | 122.0 | 4,485,511,780 |
| female 3 (F3) | 40,338,972 | 120.9 | 4,876,981,715 |
| female 4 (F4) | 37,106,658 | 120.8 | 4,482,484,286 |
| female 5 (F5) | 33,837,050 | 122.1 | 4,131,938,909 |
| female 6 (F6) | 33,285,232 | 122.3 | 4,070,993,215 |
| female 7 (F7) | 33,078,934 | 122.5 | 4,051,046,244 |

#### Table S24. Expression levels of 31 DEGs in the GSD among males, females and pseudo-males

| Patterns | Gene | symbol | FPKM in males | FPKM in pseudo-males | FPKM in females |
| --- | --- | --- | --- | --- | --- |
| Higher expression in both pseudo-males and males | JMU_LC_G_21511 |  | 26.35 | 33.24 | 0.46 |
|  | JMU_LC_G_02406 | *tbx1* | 0.72 | 1.63 | 0.00 |
|  | JMU_LC_G_02404 | *tet3* | 4.30 | 1.64 | 0.08 |
|  | JMU_LC_G_00757 |  | 183.25 | 203.75 | 16.33 |
|  | JMU_LC_G_01180 |  | 4.46 | 5.10 | 0.73 |
|  | JMU_LC_G_00776 | *parp4* | 2.19 | 3.71 | 0.26 |
|  | JMU_LC_G_02419 | *arvcf* | 5.07 | 15.72 | 0.98 |
|  | JMU_LC_G_02432 | *auts2* | 2.44 | 5.67 | 0.64 |
|  | JMU_LC_G_00765 | *wiz* | 1.45 | 3.67 | 0.28 |
|  | JMU_LC_G_02451 | *kat6a* | 13.32 | 16.04 | 2.98 |
|  | JMU_LC_G_00748 |  | 1.65 | 1.77 | 0.08 |
|  | JMU_LC_G_00749 | *pdxdc1* | 5.69 | 8.68 | 0.86 |
|  | JMU_LC_G_00767 |  | 22.17 | 34.52 | 5.77 |
|  | JMU_LC_G_02420 | *dmrt1* | 2.12 | 8.73 | 0 |
|  | JMU_LC_G_18270 | *dmrt2* | 0.043 | 0.173 | 0 |
|  | JMU_LC_G_02588 | *dmrt3* | 0.14 | 0.03 | 0 |
|  | JMU_LC_G_02662 | *smarca2* | 16.49 | 14.28 | 2.30 |
|  | JMU_LC_G_01404 | *npffr2* | 0.083 | 0.19 | 0 |
| Higher expression in females | JMU_LC_G_02421 | *rnf* | 3.77 | 6.74 | 119.31 |
|  | JMU_LC_G_02408 | *flo11* | 2.45 | 4.86 | 45.07 |
|  | JMU_LC_G_02405 | *gnb1l* | 4.95 | 8.01 | 484.79 |
|  | JMU_LC_G_02410 | *pgam5* | 6.28 | 20.23 | 433.23 |
|  | JMU_LC_G_02417 | *zdhhc8* | 0.77 | 1.35 | 21.68 |
|  | JMU_LC_G_02418 | *chfr* | 10.47 | 18.95 | 95.16 |
|  | JMU_LC_G_00747 | *trappc5* | 3.54 | 6.25 | 54.52 |
|  | JMU_LC_G_00768 | *ttf2* | 10.22 | 19.35 | 157.48 |
|  | JMU_LC_G_01175 | *dtd1* | 3.49 | 11.15 | 121.19 |
|  | JMU_LC_G_02593 | *fbp2* | 1.60 | 4.44 | 39.79 |
|  | JMU_LC_G_10237 | *fbp1* | 0.42 | 1 | 20.78 |
|  | JMU_LC_G_02604 | *kank1* | 1.82 | 5.16 | 20.02 |

#### Table S25. Primers of two DEGs for qRT-PCR analysis.

| **Gene Symbol** | **Gene ID** | **Primer sequence (5′-3′)** | **Annealing temperature** |
| --- | --- | --- | --- |
| *rnf* | JMU_LC_G_02421 | F: CCCTTCAGTCTCGGTGCTTAG | 60℃ |
|  |  | R:CAGGCAGAACGTGTGCTTG |  |
| *dmrt1* | JMU_LC_G_02420 | F:TTGCTCCAGGAAGTCGCTC | 60 ℃ |
|  |  | R:GTGGGGCATTTTCTGATATTG |  |
| *β-actin* |  | F:TTATGAAGGCTATGCCCTGCC | 60 ℃ |
|  |  | R:TGAAGGAGTAGCCACGCTCTGT |  |

#### Table S26. The functions of 66 genes significantly co-expressed with *dmrt1*.

(in a separate excel file)

#### Table S27. GO and KEGG pathway enrichment for hormone-induce DEGs

(in a separate excel file)

#### Table S28. GO and KEGG pathway enrichment for genetic-determining specific DEGs

(in a separate excel file)

#### Table S29. The GO and KEGG pathways by common DEGs shared in both GSD and hormone-induced ESR

(in a separate excel file)
